# Genetic Drivers of Sensitivity or Resistance to RAS(ON) Multi-Selective Inhibitors in *NRAS*-Mutated Melanoma

**DOI:** 10.64898/2025.12.18.695203

**Authors:** Mona Foth, Wontak Kim, Kayla O’Toole, Brandon Murphy, Montserrat Justo-Garrido, Sanjana Boggaram, Phaedra Ghazi, Euan Brennan, M. Isaac Wright, Tate Shepherd, Emilio Cortes Sanchez, Yingyun Wang, Jennifer A. Roth, Matthew Rees, Melissa Ronan, Jingjing Jiang, Urszula Wasko, Amanda Jiang, Carly Becker, Dekker C. Deacon, Siwen Hu-Lieskovan, Conan Kinsey, Jeffery Russell, Aparna Hegde, Ignacio Garrido-Laguna, Matthew Holderfield, Mallika Singh, Martin McMahon

## Abstract

Most patients with advanced *BRAF* or *NRAS*-driven melanoma receive front-line immunotherapy. However, if immunotherapy fails, *BRAF*-mutated patients have effective second-line therapies, whereas *NRAS*-mutated patients lack pathway-targeted options. Recently, RAS(ON) multi-selective inhibitors like RMC-7977, and the investigational agent daraxonrasib, were described that, in partnership with cyclophilin-A (CYPA), inhibit RAS[GTP] signaling. Both compounds demonstrate potent anti-proliferative activity against *NRAS*-mutated melanoma cell lines and robust anti-tumor activity against preclinical melanoma models. However, in preclinical models, resistance to RMC-7977 monotherapy arose through mutations in *Ppia* (encoding CYPA) or *Map2k1* (encoding MEK1). Moreover, two clinical case studies in patients with *NRAS*-mutated melanoma treated with daraxonrasib demonstrated clear anti-tumor activity in one patient, but progressive disease in another with co-occurring *NRAS* and *MAP2K1* mutations at baseline. These findings support the potential for daraxonrasib in treatment of patients with *NRAS*-mutated melanoma, and reveal candidate mechanisms of monotherapy resistance, underscoring the need for combination therapies to improve outcomes.

**SIGNIFICANCE:** There are no pathway-targeted therapies for patients with *NRAS*-mutated melanoma. Here we demonstrate that direct pharmacological inhibition of RAS[GTP] with RMC-7977 or daraxonrasib (RMC-6236) has profound inhibitory effects in preclinical models of *NRAS*-mutated melanoma. Furthermore, we identify mechanisms of resistance to RMC-7977 through mutational inactivation of CYPA or mutational activation of MEK1.

## INTRODUCTION

Melanoma is the sixth most common cancer diagnosed in the U.S. and the deadliest form of skin cancer. Mutationally-activated NRAS is expressed in ∼25% of all human melanomas, with most mutations occurring at codon 61, or less frequently at codons 12 or 13 (1,2). Such mutationally-altered oncoproteins (e.g., NRAS^Q61R^, NRAS^G12V^, NRAS^G13D^) exhibit both loss of GTPase function and enhanced signaling function (3). NRAS oncoproteins are reported to activate both RAF protein kinases and PI3’-lipid kinases leading to downstream activation of RAF>MEK>ERK MAP kinase and PI3’-lipid signaling, respectively (4). Although the coordinate activation of these kinase effector pathways is central to melanoma initiation and progression, and is required for multiple aberrant behaviors of melanoma cells, combinatorial efforts to pharmacologically target these pathways in NRAS-driven melanoma patients have resulted either in unacceptable toxicity, lack of efficacy, or both (5–7). Almost all patients with advanced melanoma, regardless of genetic driver alterations, receive immunomodulatory agents as first-line therapy. However, for those patients with *NRAS*-mutated melanoma who are either ineligible for immunotherapy, or for whom immunotherapy fails, second-line treatment options are limited. Despite deep knowledge of key signaling pathways downstream of NRAS[GTP], there are no FDA-approved pathway-targeted therapies for treating NRAS-driven melanoma.

Despite the prevalence of *RAS* mutations in cancer (∼20%), and longstanding interest in direct pharmacological targeting of RAS oncoproteins for cancer therapy, the development of such inhibitors has proven challenging (2,8). However, the proof-of-concept development of covalent inhibitors of KRAS^G12C^ spurred a renaissance in efforts to pharmacologically target RAS oncoproteins directly. Indeed, inhibitors of RAS[GTP], RMC-7977 (tool compound) and daraxonrasib (RMC-6236; investigational agent) have been developed as multi-selective inhibitors of both the mutationally-activated and the normal forms of H-, K– or NRAS proteins (9,10). Upon entering the cell, these agents bind to cyclophilin A (CYPA), an abundant cytosolic cis-trans peptidyl-prolyl isomerase believed to serve as a chaperone for intracellular protein folding. Then the RMC-7977⇔CYPA binary complex binds directly to RAS[GTP]. Binding of the binary complex to RAS[GTP] inhibits RAS oncoprotein signaling by two mechanisms that are independent of CYPA’s cis-trans peptidyl-prolyl isomerase activity: 1. Promotes steric displacement of downstream effectors such as RAF and PI3’-kinase from RAS[GTP]; and 2. For certain RAS oncoproteins (e.g., KRAS^G12D^) complex formation promotes hydrolysis of the RAS oncoprotein-bound GTP thereby converting RAS back to its signaling inactive, GDP-bound state (9,11–14). Notably, RMC-7977 and daraxonrasib do not promote GTP hydrolysis of KRAS^Q61H^ or KRAS^Q61L^ oncoproteins (13). Both agents display potent anti-cancer activity against RAS-transformed cells *in vitro* and in numerous preclinical models, leading to the advancement of daraxonrasib into clinical trials, e.g., NCT05379985, NCT06625320, NCT06445062, NCT06162221 (9,14,15). Indeed, initial clinical data have shown encouraging efficacy and tolerability profiles, supporting further investigation (10). RAS(ON) multi-selective inhibitors such as RMC-7977 and daraxonrasib are not selective solely for mutationally-activated RAS oncoproteins, which would explain the activity of these compounds against cancers that rely on wild-type RAS signaling for their maintenance such as those driven by mutationally-activated receptor tyrosine kinases or by silencing of the neurofibromatosis 1 (*NF1*) gene that encodes a RAS GTPase-activating protein (9).

While single-agent anti-RAS therapies hold promise for the treatment of NRAS-driven melanoma, the emergence of acquired drug resistance that limits the durability of patients’ responses remains a significant challenge (16–18). Several putative resistance mechanisms for RAS-selective treatments – both genetic and non-genetic –have been described in the literature: 1. On-target alterations that render the RAS oncoprotein drug resistant or over-expressed; 2. RAS-independent activation of downstream RAF>MEK>ERK or PI3’-kinase signaling; 3. Activation of bypass pathways such as YAP/TAZ>TEAD signaling (18,19); or 4. Cell plasticity as well as transcriptional or histopathologic changes (20–22). Such pleiotropic mechanisms of resistance have emerged even within a single KRAS^G12C^-driven lung cancer patient following treatment with adagrasib (17). Previous work on RMC-7977 resistance in preclinical models of pancreatic cancer revealed elevated expression of c-MYC in resistant tumors (15). In addition, epigenetic changes and adaptive rewiring of the tumor microenvironment, such as increased availability of growth factors may contribute to drug resistance by creating a more supportive niche for cancer cells under therapeutic pressure (8). Finally, intra-tumoral heterogeneity and clonal evolution contribute to the emergence of drug-resistant sub-populations, further complicating therapeutic interventions (8,16).

Here we observed that preclinical models of NRAS-driven melanoma were generally exquisitely sensitive to the anti-tumor effects of RMC-7977, whereas BRAF^V600E^-driven melanoma models were resistant, as anticipated. However, some of the NRAS-driven melanoma models displayed emergence of drug resistance that was due to either: 1. Mutations silencing CYPA expression or; 2. Mutations that led to RAS-independent, constitutive activation of MEK1>ERK signaling. While these data confirm the essentiality of CYPA in the mechanism of action of RMC-7977, and the primacy of RAF>MEK>ERK signaling in the maintenance of NRAS-driven melanoma, they do not explain why MEK1/2 inhibitors have little or no clinical activity in this disease. Moreover, such multi-faceted resistance strategies underscore the complexity of targeting the RAS pathway and highlight the need for novel combination therapies to overcome these challenges.

## RESULTS

### *NRAS* mutation predicts for RMC-7977 sensitivity in pan-cancer analysis of 573 cell lines

To identify genetic alterations associated with RMC-7977 sensitivity or resistance, the anti-proliferative potency of RMC-7977 was assessed in a large panel of human cancer cell lines **(Fig. 1A),** as well as selectively in melanoma cell lines **(Fig. 1B),** in a multiplexed format (PRISM assay; **Suppl. Table S1).** Consistent with published data (2,3), cell lines with KRAS^G12X^ alterations (KRAS^G12X^; n=92 cell lines) displayed the highest sensitivity to RMC-7977 **(Fig. 1A).** Importantly, *NRAS*-mutated (NRAS^MUT^; n=36 cell lines) cells were the second most sensitive group **(Fig. 1A).** RAS-Pathway^Other^ (n=125 cell lines) and RAS-pathway^WT^ (n=320 cell lines) were significantly less sensitive to RMC-7977 than *KRAS*– or *NRAS*-mutated cancer cell lines (Mann-Whitney test: NRAS^MUT^ vs. RAS-Pathway^Other^ p<0.0001; NRAS^MUT^ vs. RAS-pathway^WT^ p<0.0001; NRAS^MUT^ vs. KRAS^G12X^ p=ns, **Fig. 1A**). When we selectively examined only melanoma cell lines by PRISM assay, those with *NRAS* mutations displayed the highest sensitivity to RMC-7977 (n=7 cell lines), whereas cells driven by BRAF^V600E^ were substantially resistant (n=32 cell lines; p=0.0001; **Fig. 1B**). Finally, cells expressing other genetic drivers that do not result in NRAS^Q61X^ or BRAF^V600E^ (NRAS/BRAF^WT^; n=3 cell lines) presented as intermediate responders to RMC-7977 (Mann-Whitney test: NRAS^MUT^ vs. BRAF^MUT^ p=0.0001; BRAF^MUT^ vs. NRAS/BRAF^WT^ p=0.0002; NRAS^MUT^ vs. NRAS/BRAF^WT^ p=ns, **Fig. 1B**).

**FIGURE 1:**
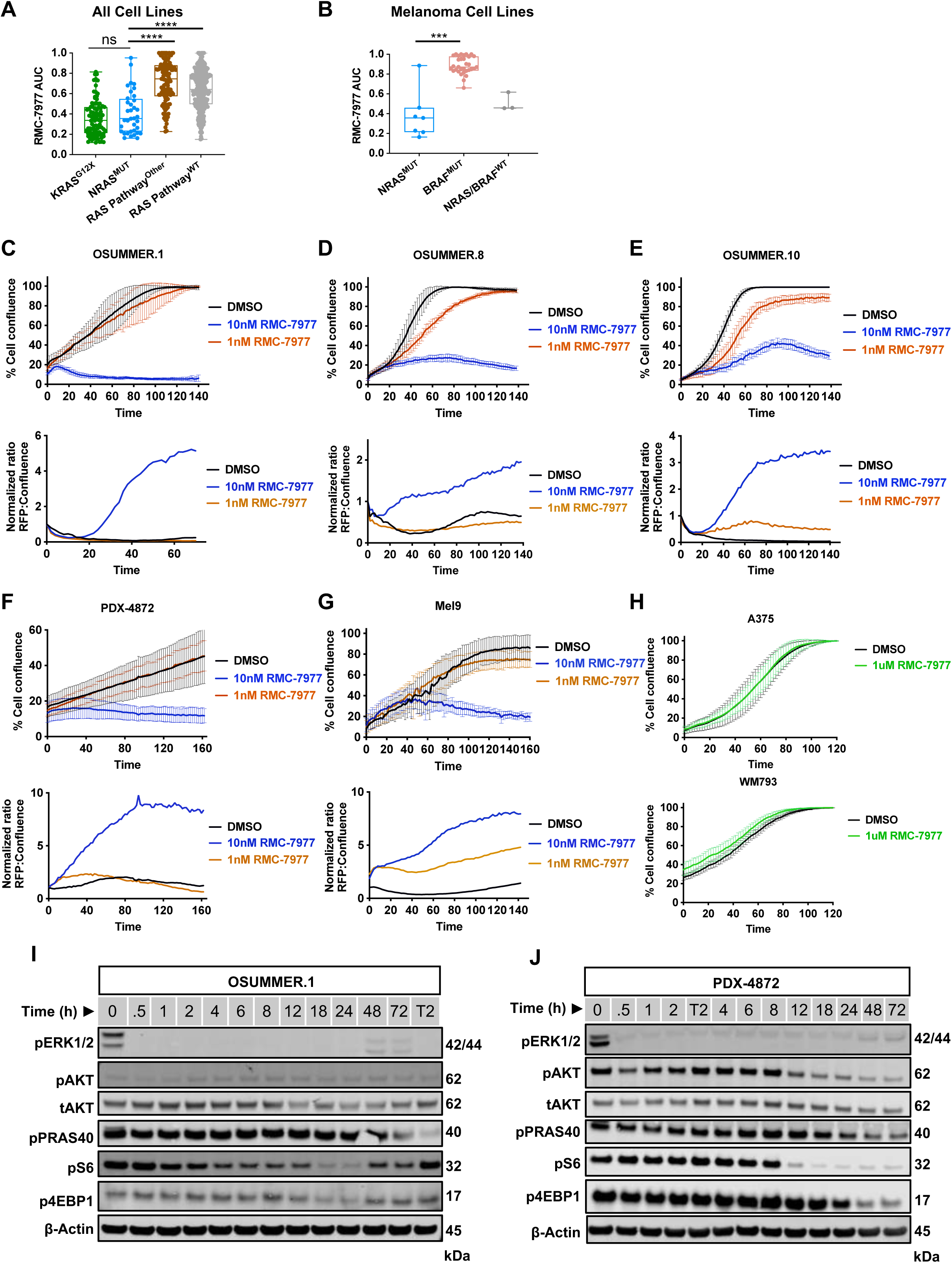
Potent and selective anti-proliferative activity of RMC-7977 against *NRAS*-mutated melanoma cells. **(A-B)** Cell viability assay of n=573 human tumor cell lines **(A)** or selectively of melanoma cell lines **(B)** in a pooled, multiplexed format (PRISM assay). Area under the curve (AUC) difference between cell lines by genotype as indicated. Points represent AUC for individual cell lines. Low AUC values indicate sensitivity; high AUC values indicate resistance. Mann-Whitney test: ***, p = 0.0001; ****, p < 0.0001, ns, non-significant. **(C-H)** Cell confluency or cytotoxicity assays under pharmacological inhibition *in vitro*. Mouse NRAS-driven melanoma cell lines OSUMMER.1 (NRAS^Q61R^, **C**), OSUMMER.8 (NRAS^Q61K^, **D**), or OSUMMER.10 (NRAS^G13R^, **E**), or human NRAS-driven melanoma cell lines PDX-4872 (NRAS^Q61K^, **F**), Mel9 (NRAS^Q61R^, **G**), or human BRAF^V600E^-driven melanoma cell lines A375 and WM793 (**H**) were treated with vehicle (DMSO) or RAS(ON) multi-selective inhibitor RMC-7977 at indicated concentrations and analyzed for cell confluence or cytotoxicity by CytoxRed reagent uptake using an Incucyte Live-Cell Analysis System over time. Graphs are representative of n=3 independent experiments. Error bars represent standard deviation (SD) of triplicate technical replicates. **(I-J)** Immunoblot analysis of cell lysates prepared from mouse melanoma cell lines OSUMMER.1 **(I)** or human-derived PDX-4872 **(J)** treated with DMSO or 80nM RMC-7977 over a time course of 0.5-72 hours. A two-hour 100nM trametinib control treatment (T2) was included in each run for complete suppression of pERK1/2. Lysates were analyzed for the phosphorylation (p) or total (t) abundance of ERK1/2, AKT, PRAS40, S6 ribosomal protein, 4EBP1, or β-actin, as indicated. Immunoblot quantification for **I-J** in **Supplementary Fig. S1 F-I.**

### Potent and selective anti-proliferative activity of RAS(ON) multi-selective inhibitor RMC-7977 against *NRAS*-mutated melanoma cells

A panel of mouse or human *NRAS*-mutated melanoma cell lines **(Suppl. Table S2)** derived from either suitably manipulated genetically engineered mice (OSUMMER), or were newly derived from human melanoma patient-derived xenograft (MPDX: Mel9 & PDX-4872) models generated at Huntsman Cancer Institute (HCI) were treated with RMC-7977 (23). As controls, we used two BRAF^V600E^-driven melanoma cell lines, A375 and WM793. Melanoma cells were exposed to different concentrations of RMC-7977 *in vitro* with cell proliferation or cell death assessed using Incucyte live-cell imaging and analysis **(Figs. 1C-H)**. Cell proliferation was potently inhibited in all seven NRAS-mutated melanoma cell lines **(Figs. 1C-G, Suppl. Figs. S1A-C)** by as little as 10nM RMC-7977, and completely by concentrations of ≥10nM. By contrast, both BRAF^V600E^-driven melanoma cell lines were insensitive to RMC-7977 **(Fig. 1H).** Assessment of RMC-7977-mediated cell killing (Cytotox Red) revealed increased cell death in all NRAS-driven melanoma cells at concentrations ≥10nM (**Figs. 1C-G, Suppl. Figs. S1C).**

To investigate the ability of RMC-7977 to inhibit RAS signaling, OSUMMER.1 (O.1), OSUMMER.10 (O.10), PDX-4872 or Mel9 cells were treated with RMC-7977 (80nM) over a time course of 0.5-72 hours at which time cell lysates were prepared for analysis by immunoblotting. Blots were probed with backbone– or phospho-specific antisera as indicated **(Fig. 1 I-J, Suppl. Fig. S1 D-I)**. All cells treated with RMC-7977 displayed a rapid decrease in pERK1/2 (p-T202/Y204) that was detected within 30 minutes after compound addition **(Figs. 1 I-J, Suppl. Figs. S1 D-I)**. In O.1 and O.10 cells, there was modest reactivation of pERK1/2 48 hours after RMC-7977 addition **(Fig. 1 I; Suppl. Fig. S1D, F-G),** while in PDX-4872 and Mel9 cells, pERK1/2 remained inhibited for up to 72 hours **(Fig. 1 J; Suppl. Fig. S1E, H-I).** Interestingly, there was little or no decrease in pAKT (pS473 – a surrogate for PI3’-lipid signaling) in O.1 or O.10 cells at any time point analyzed, which was consistent with little or no change in pPRAS40 (pT246), a known AKT substrate. However, decreased pAKT was observed in human PDX-4872 and Mel9 cells albeit at later time points >8 hours after RMC-7977 addition. All cells displayed decreased pRP-S6 (pS235/236) at later times after RMC-7977 addition: >4 hours in O.1 cells, >8 hours in O.10, PDX-4872, and Mel9 cells. Finally, we noted decreased phosphorylation of 4E-BP1 (pS65) in all cells, but with delayed kinetics compared to changes in pRP-S6 **(Fig. 1 I-J, Suppl. Fig. S1 D-I)**. These data are consistent with the RAF>MEK>ERK MAP kinase signaling pathway being a primary conduit of intracellular NRAS oncoprotein signaling (9,10,15). However, the inconsistent changes in pAKT and downstream outputs from mTORC1/2 such as pPRAS40, pRP-S6 and p4E-BP1 that are also reported to be downstream of PI3’-lipid signaling, might suggest that there are other factors influencing this signaling axis in NRAS-driven melanoma cells. Although we used 80nM RMC-7977 for these biochemical assays to approach clinically relevant exposure levels to daraxonrasib, lower concentrations of RMC-7977 (25nM: ∼2× GI99 in O.1) were sufficient to achieve comparable inhibition of pERK1/2. Effects on the PI3K/AKT pathway remained minimal at either concentration **(Suppl. Fig. S2A-D).**

### Inhibition of NRAS[GTP] signaling displays robust anti-tumor activity against NRAS-driven melanoma models

To test the anti-tumor activity of RMC-7977 against preclinical NRAS-driven mouse models of melanoma, we first evaluated the agent’s pharmacodynamics using O.1 allografts. Tumor bearing immunocompetent mice were administered a single dose of RMC-7977 (10 mg/kg) followed by analysis of tumor lysates by immunoblotting or by immunohistochemical analysis of formalin-fixed sections at 6, 8, or 24 hours post compound administration. We observed profound inhibition of pERK1/2 at 6 and 8 hours post-treatment, followed by full recovery at 24 hours **(Suppl. Fig. S3A-B)**. However, consistent with prior immunoblot analysis of O.1 cells, we observed no modulation of pAKT in these tumor samples (Suppl. Fig. S3C-E).

Next, the responses to RMC-7977 of O.1, O.8 or O.10 allografted tumors were monitored over 24 days in immunocompetent *rGH-ffLuc2-eGFP* (aka *Glowing Head, GH*) mice (24). Tumors were treated with RMC-7977 (10 mg/kg, qd po) when they were ∼300 mm^3^. A robust anti-tumor response to RMC-7977 was observed in all three models, indicated by a marked reduction in tumor burden during the first 10 days of treatment **(Figs. 2A-C).** Furthermore, even large (600-1000 mm^3^) O.8– or O.10-derived tumors in two mice that had surpassed the regular enrollment window for treatment (250-350 mm^3^) responded to RMC-7977 **(Suppl. Fig. S3F, G)**. Both such mice exhibited striking responses within a few days of treatment, leading to a marked reduction in tumor volume **(Suppl. Fig. S3F, G)**. However, as described below, RMC-7977 resistant O.1 and O.10 tumors emerged within 14-21 days of treatment. In parallel, we harvested control vs. treated (2 or 24 days) O.1, O.8 or O.10 tumors for analysis of pERK1/2 by immunohistochemistry **(Fig. 2D)**. Vehicle control treated tumors displayed readily detectable pERK1/2, which was potently suppressed following two days of RMC-7977 treatment. O.8 tumors still had diminished pERK1/2 at 24 days after RMC-7977 treatment. By contrast, pERK1/2 was restored in O.1 and O.10 tumors after 24 days of treatment, when resistant tumors were close to endpoint.

**FIGURE 2:**
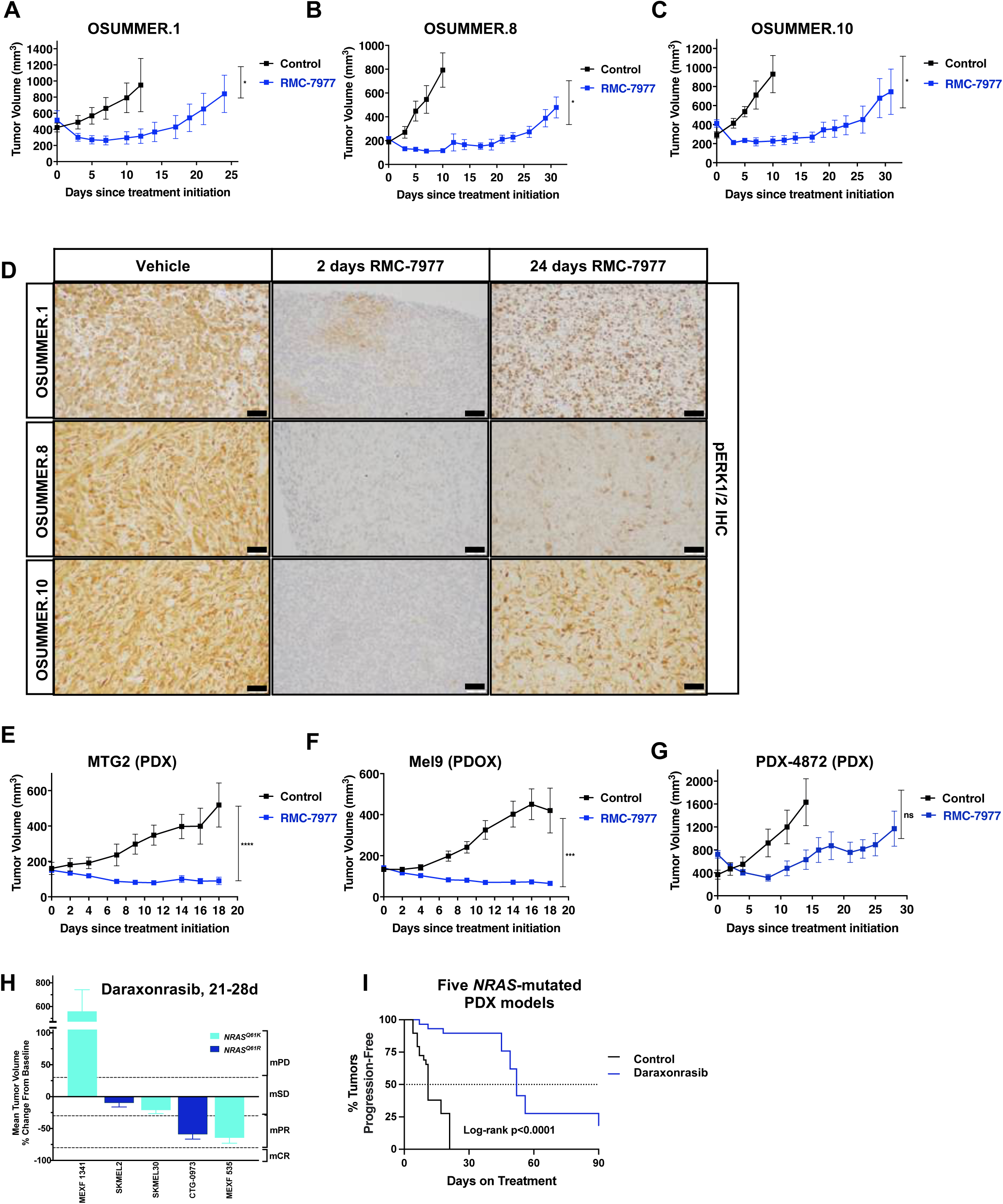
Inhibition of NRAS[GTP] signaling displays robust anti-tumor effects against NRAS-driven melanoma models. **(A-C)** Tumor volume graphs of allografted OSUMMER cells in immunocompetent Glowing Head mice treated with vehicle control (Veh) or 10 mg/kg RMC-7977 (RMC) over 24-31 days. OSUMMER.1 (NRAS^Q61R^; Veh n=6 vs. RMC n=6; p=0.0477; **A**), OSUMMER.8 (NRAS^Q61K^, Veh n=6 vs. RMC n=7; p=0.00194; **B**), or OSUMMER.10 (NRAS^G13R^, Veh n=6 vs. RMC n=6; p=0.0437; **C**) Error bars represent SEM. (Mann-Whitney test: ns, non-significant; *, p < 0.05; **, P < 0.01; ***, P < 0.001; ****, P < 0.0001). **(D)** Representative sections of immunohistochemistry (IHC) for pERK1/2 on OSUMMER.1, OSUMMER.8, or OSUMMER.10 allograft tumors of mice that were treated with vehicle or 10 mg/kg RMC-7977 for 2 or 24 days as indicated. Scale bar in lower right corner represents 50uM in all images. **(E-G)** Tumor volume graphs of patient-derived xenografts in immunocompromised NOD/SCID mice treated with vehicle or 10 mg/kg RMC-7977 as indicated over 18-28 days. MTG2 (NRAS^Q61R^, Veh n=4 vs. RMC n=6; p<0.0001; **E**), patient-derived organoid xenograft (PDOX) Mel9 (NRAS^Q61R^, Veh n=5 vs. RMC n=6; p=0.0002; **F**) or patient-derived xenograft PDX-4872 (NRAS^Q61K^, Veh n=7 vs. RMC n=6; p=0.6165; **G**). Error bars represent SEM. (Mann-Whitney test: ns, non-significant; *, p < 0.05; **, P < 0.01; ***, P < 0.001; ****, P < 0.0001). **(H-I)** Five PDX mouse melanoma models driven by NRAS^Q61K^ or NRAS^Q61R^ treated with vehicle (n=29) or 25 mg/kg daraxonrasib (n=29) over 90 days. **(H)** Mean tumor volume % change from baseline of five NRAS-driven models treated with daraxonrasib at 25 mg/kg as indicated. NRAS^Q61K^ (teal) or NRAS^Q61R^ (blue). Tumor volume change by mRECIST criteria; mPD: Progressive disease, mSD: Stable disease, mPR: Partial response, mCR: Complete response. Error bars represent SEM. **(I)** Tumor progression-free survival of the five PDX models. Median progression-free survival was 11 days (vehicle control) vs. 52 days (daraxonrasib); p<0.0001 Log Rank (Mantel-Cox) test.

We next tested the anti-tumor effects of RMC-7977 against three human MPDX models: MTG2 (NRAS^Q61R^); Mel9 (NRAS^Q61R^); and PDX-4872 (NRAS^Q61K^) xenografted into NOD/SCID mice (Suppl. Table S3). Similar to OSUMMER allografts, RMC-7977 demonstrated robust anti-tumor activity against all three of these models **(Figs. 2E-G)**. Mice bearing MTG2 or Mel9 MPDXs displayed tumor regression up to 18 days of treatment **(Figs. 2E-F)**, whereas mice bearing PDX-4872 tumors displayed tumor regression during the first 7 days of treatment **(Fig. 2G)**. However, in this MPDX model, we observed emergence of on-treatment resistance after 7-10 days of treatment. At euthanasia, analysis of H&E stained sections revealed that RMC-7977-treated Mel9 tumors displayed a hyalinizing stroma, a treatment effect that is observed in human tumor specimens after effective therapy **(Suppl. Fig. S3H).**

In parallel to the analysis described above, we tested the sensitivity of five NRAS-driven melanoma models (MEXF-1341, SK-MEL2, SK-MEL30, CTG-0973 and MEXF-535) to the clinical investigational agent, daraxonrasib **(Figs. 2H-I).** Daraxonrasib treatment (25 mg/kg, qd, po) elicited partial responses in two MPDX models (CTG-0973 & MEXF-535), stable disease in two cell line models (SK-MEL2 & SK-MEL30) and the fifth MPDX model (MEXF-1341) displayed progressive growth **(Fig. 2H)**. Overall, daraxonrasib significantly prolonged time to tumor doubling (a surrogate for progression-free survival) in these models tested (52 days vs. 11 days; p<0.0001) **(Fig. 2I).**

To further characterize RAS pathway modulation by daraxonrasib *in vivo*, we carried out pharmacodynamic (PD) assessments in the SK-MEL30 (NRAS^Q61K^) xenograft model, either following a single dose or at day 45 following tumor progression on treatment **(Suppl. Fig. S3I).** The 25 mg/kg qd po dose of daraxonrasib was shown via translational modeling to achieve >90% RAS pathway suppression over a 24-hour dosing interval in xenograft models dependent on RAS signaling for growth, consistent with previously published data (10). This dose is considered equivalent to approximately 300mg daily in humans, the dose and regimen selected for the ongoing Phase 3 RASolute 302 trial (NCT06625320). NRAS>RAF>MEK>ERK pathway modulation was assessed by measuring human DUSP6 mRNA expression levels, which revealed profound DUSP6 mRNA suppression at 4 and 8 hours following a single 25 mg/kg dose of daraxonrasib treatment, consistent with strong target engagement. In contrast, tumors collected at the end of the study, during apparent on-treatment relapse, exhibited evidence of RAS pathway reactivation **(Suppl. Fig. S3I).**

### RMC-7977 elicits robust anti-tumor activity against autochthonous NRAS-driven melanomas

To complement experiments in transplantation models, we employed genetically engineered mouse (GEM) models of melanoma. The first, *<u>T</u>yr::CreER^T2^*; *<u>N</u>ras^LSL-Q61R/Q61R^; <u>C</u>dkn2a^flox/flox^* (*TNC,* **Suppl. Fig. S4A**) is driven by melanocyte-specific expression of NRAS^Q61R^ combined with silencing of the *Cdkn2a* locus encoding the p16^INK4A^ and p19^ARF^ tumor suppressors (25–28). The second, *Tyr::CreER^T2^; Braf^CA/+^; Cdkn2a^flox/flox^* (*TBC***, Suppl. Fig. S4B**) is driven by BRAF^V600E^ expression also combined with INK4A/ARF silencing. Topical application of 4-HT to a single location on the back skin of adult (≥6 weeks) *TNC* or *TBC* mice, was followed by exposure to a single dose of 2000-3000J/m^2^ of UVB irradiation **(Suppl. Fig. S4C-D)**, which accelerates melanomagenesis in GEM models (29,30). Under these conditions, melanomagenesis was observed in adult *TNC* mice at a median of 70 days (2000J/m^2^ UVB) or 85 days (3000J/m^2^ UVB) post-initiation (Log Rank, p=ns). 4-HT induction without UVB-irradiation resulted in a median melanoma onset of 140 days in *TNC* mice (no UVB vs. 3000J/m^2^ UVB, Log Rank p=0.0128, **Fig. 3A**). Comparable in their melanoma onset, *TBC* mice induced as adults followed by a single 3000J/m^2^ UVB exposure developed melanomas at a median onset of 52 days post-initiation (TBC 3000J/m^2^ vs. TNC 3000J/m^2^, Log Rank p=ns; **Fig. 3B).** NRAS^Q61R^-driven tumors exhibited typical markers of mouse melanoma, including S100A4 expression and markers of proliferation such as Ki67, and detectable pERK1/2 as an immunohistochemical indicator of RAF>MEK>ERK (pERK1/2) signaling **(Suppl. Fig. S4E).** By contrast, there was barely detectable pAKT or pPRAS40 as markers of PI3’-lipid signaling, although pRPS6 was detected by immunoblotting **(Suppl. Fig. S4F).** It should be noted that application of 4-HT to neonatal *TNC* mice **(Suppl. Fig. S4C)** also led to the development of multiple melanomas across their skin most likely due to maternal grooming, which presented a technical challenge for consistent tumor measurement and treatment stratification.

**FIGURE 3:**
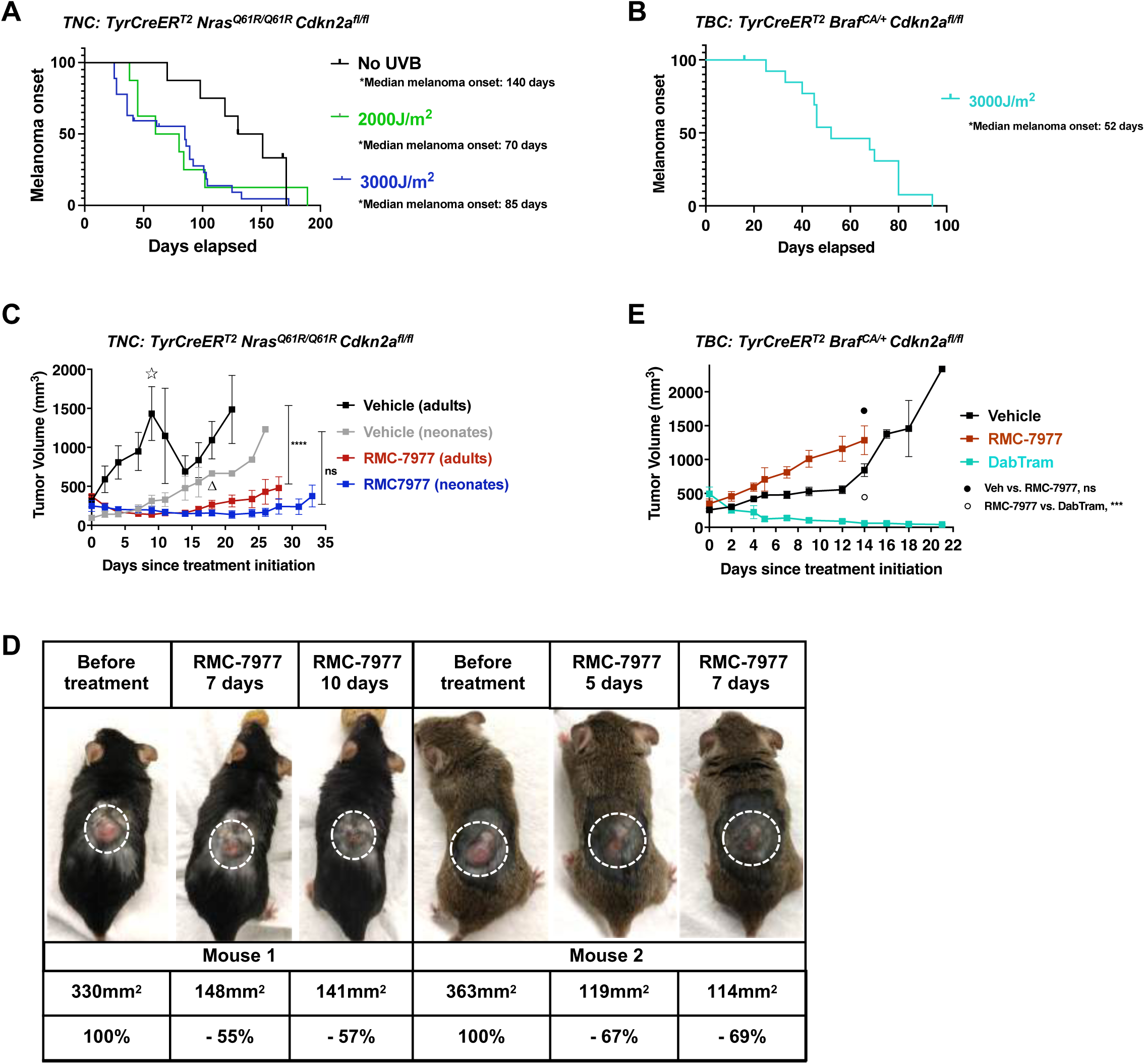
RMC-7977 elicits robust anti-tumor effects against autochthonous NRAS-driven melanomas. (**A)** Kaplan-Meier plot of melanoma onset following 4-HT induction and UV-irradiation of adult *TNC* mice at indicated UVB doses. Median melanoma onset in *TNC* model was 70 days (2000J/m^2^; n=8) or 85 days (3000J/m^2^; n=24; Log Rank (Mantel-Cox) p=ns compared to 2000J/m^2^). Melanoma onset following 4-HT induction without UV-irradiation was 140 days (n=8; no UVB vs. 3000J/m^2^; p=0.0128). **(B)** Kaplan-Meier plot of melanoma onset following 4-HT induction and UV-irradiation of adult *TBC* mice at 3000J/m^2^. Median melanoma onset in *TBC* model was 52 days (n=13; Log Rank (Mantel-Cox) p=ns compared to *TNC* induced at 3000J/m^2^). **(C)** Tumor volume graph of *TNC* mice induced as neonates or adults, and treated with vehicle or 10 mg/kg RMC-7977 as indicated (vehicle adults n=6; vehicle neonates n=6; RMC-7977 adults n=4; RMC-7977 neonates n=9). Each dot represents the average tumor volume per mouse, i.e. one tumor in mice induced as adults, or the average of multiple tumors per mouse per time point in mice induced as neonates. Star (11) above adult-vehicle group on day 9 indicates time point where n=3 mice had to be sacrificed and removed from the experiment due to tumor sizes reaching end point, which resulted in a drop of the average tumor volume of the remaining n=3 mice in the adult-vehicle treated group. Triangle (∇) in the neonates-vehicle group on day 18 indicates the end point of five mice due to the size and location of the tumors, where only one was continued from day 18 to day 26 in the vehicle treated group. Error bars represent SEM. (Mann-Whitney test: Adult-vehicle vs. Adult-RMC-7977 at 33 days endpoint p < 0.0001, ****; Neonates-vehicle vs. Neonates-RMC-7977 sample number too small at 33 days endpoint, p=0.0516, ns; or at day 16 (p=0.5737). **(D)** Representative images of two *TNC* mice induced as adults before treatment and during early tumor regression on RMC-7977 on treatment days 7 and 10, or days 5 and 7, respectively. White dotted circle indicates location of tumor. Percent reduction in tumor volume as indicated below photographs. **(E)** Tumor volume graph of *TBC* mice induced as adults and treated with vehicle, 10 mg/kg RMC-7977, or dabrafenib+trametinib (Dab/Tram; 30 mg/kg Dab; 1 mg/kg Tram) as indicated (vehicle n=3; RMC-7977 n=5; DabTram n=4). Error bars represent SEM. Mann-Whitney test at day 14: Vehicle vs. RMC-7977 p=0.0650, ns, black dot; RMC-7977 vs. DabTram p=0.0006, ***, white circle).

To test the anti-tumor activity of RMC-7977 in these GEM models, melanomas were initiated in either neonatal or adult *TNC* mice and, when tumors were measurable, they were treated with either RMC-7977 (10 mg/kg, qd, po) or vehicle. In *TNC* mice initiated as neonates, RMC-7977 elicited robust regression of multi-focal melanomas regardless of their location on the skin (**Fig. 3C).** Indeed, one mouse initiated as a neonate developed five individual tumors across its body, all of which responded to RMC-7977 for up to 30 days. Upon cessation of RMC-7977 administration, all five tumors displayed re-growth **(Suppl. Fig S5A)**, but all such melanomas remained treatment sensitive following re-challenge with RMC-7977 **(Suppl. Fig S5A).** Interestingly, the four tumors that were harvested for histological analysis demonstrated marked differences in their histomorphological appearance, as well as in the level of pERK1/2 detected at endpoint **(Suppl. Fig. S5B).**

RMC-7977 treatment also elicited potent anti-tumor activity against melanomas initiated in adult *TNC* mice as evidenced by significant tumor regression over ∼14 days (Mann-Whitney test, p=<0.0001; **Fig. 3C)** with some tumors decreasing by up to 70% in size within the first seven days of treatment **(Fig. 3D)**. By contrast, RMC-7977 had no anti-tumor effect against BRAF^V600E^-driven melanomas initiated in adult *TBC* mice (Mann-Whitney test p=0.6574, ns; **Fig. 3E).** However, as expected, BRAF^V600E^-driven melanomas arising in *TBC* mice were sensitive to dabrafenib plus trametinib (Mann-Whitney test, p=<0.0001; **Fig. 3E).** As observed in the transplantation models, we noted emergence of resistance in the *TNC* model after ≥14 days of treatment **(Fig. 3C).** Moreover, we observed re-activation of RAF>MEK>ERK signaling by immunoblotting in all RMC-7977-treated mouse tumor lysates (n=3 at end-point; **Suppl. Fig. S4F**).

Overall, preclinical testing of RMC-7977 revealed that: 1. Most NRAS-driven melanoma models are initially exquisitely sensitive to anti-tumor effects following treatment with one exception (MEXF-1341) that displayed primary resistance; 2. BRAF^V600E^-driven melanomas are resistant to RMC-7977 treatment and; 3. The emergence of RMC-7977 resistant melanomas was observed, often quite rapidly, in both transplantation and autochthonous models. Finally, as reported previously, exposure to RMC-7977 or daraxonrasib was well-tolerated in both immunocompetent and immunodeficient mice (31).

### Analysis of mechanisms of RMC-7977 resistance in NRAS-driven melanoma cells

Many of the initially sensitive preclinical models of NRAS-driven melanoma displayed the emergence of RMC-7977 resistance (RMC-7977^R^) within 7-14 days after treatment **(Figs. 2A-C; 2F, 2G & 3C)**. To determine molecular mechanisms of resistance, we excised RMC-7977^R^ melanomas, generated and plated single cells in media supplemented with escalating concentrations of RMC-7977 from 10nM to 1μM. Using these conditions, we were readily able to generate RMC-7977^R^ O.1 or O.10 melanoma-derived cell lines at all concentrations of RMC-7977 *in vitro*, indicating the tumor-cell autonomous nature of RMC-7977 resistance. Incucyte live cell imaging analysis of cell proliferation **(Fig. 4)** revealed that, while the parental OSUMMER.1 or OSUMMER.10 cells were sensitive to doses of RMC-7977 as low as 10nM, O.1 or O.10 cell lines derived from RMC-7977^R^ tumors (OSUMMER.1: O.1^RM1^, O.1^RM2^, O.1^RM5^ and O.1^RM6^; OSUMMER.10: O.10^RM1^, O.10^RM2^, O.10^RM3^, and O.10^RM4^) continued to proliferate in concentrations of RMC-7977 up to 1μM **(Fig. 4A-B).**

**FIGURE 4:**
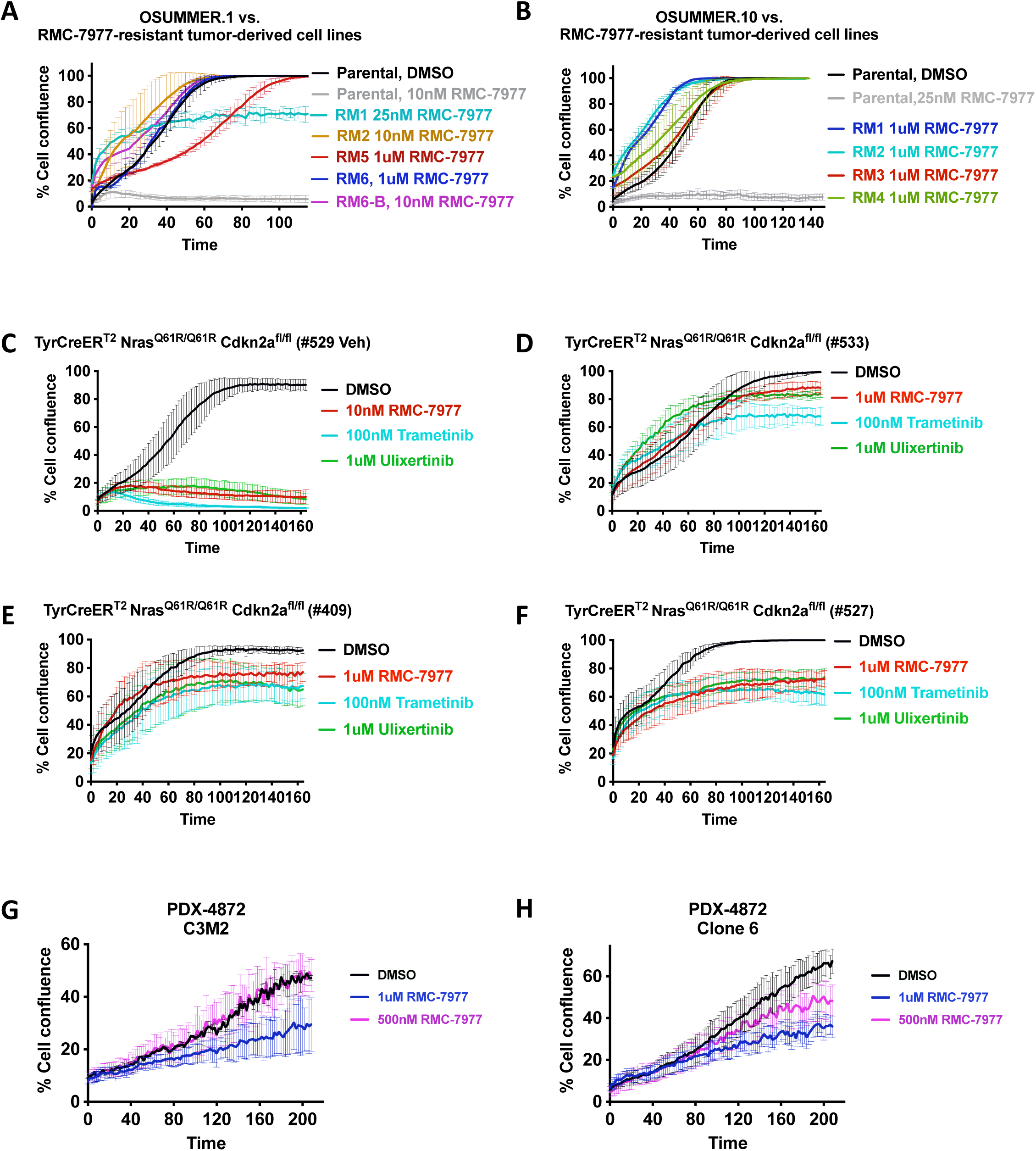
Emergence of resistance to RMC-7977 in NRAS-driven preclinical melanoma models. (**A-H)** Cell confluency assays of RMC_7977^R^ cell cultures derived from preclinical melanoma mouse models dosed with RMC-7977 until emergence of resistance. RMC-7977 resistant cell cultures were treated with vehicle (DMSO), RMC-7977, MEK1/2 inhibitor trametinib, or ERK1/2 inhibitor ulixertinib at indicated concentrations and analyzed for cell confluence using Incucyte Live-Cell Analysis. **(A)** RMC_7977^R^ cell lines derived from four RMC-7977-treated OSUMMER.1 (NRAS^Q61R^) allograft mice (O.1^RM1^, O.1^RM2^, O.1^RM5^, O.1^RM6^). Tumor cell lines O.1^RM6^ and O1^RM6-B^ were derived from the same mouse, but cultured at either 10nM (O.1^RM6-B^) or 1μM RMC-7977 (O.1^RM6^). **(B)** RMC_7977^R^ cell lines derived from four RMC-7977-treated OSUMMER.10 (NRAS^G13R^) allograft mice (O.10^RM1^, O.10^RM2^, O.10^RM3^, O.10^RM4^). **(C-F)** RMC-7977 sensitive or resistant cell lines derived from *TNC (TyrCreER^T2^; Nras^LSL-Q61R/Q61R^; Cdkn2a^flox/flox^)* mice that were dosed with either vehicle (#529-Veh) or 10 mg/kg RMC-7977 (#533, #409, and #527) daily for 28 days until resistance emerged. Tumor-derived cell lines #529, #533, #409, or #527 were prepared and cultured in the absence (#529-Veh, **C**) or presence (#533 **(D**), #409 **(E)**, or #527 **(F)**) of RMC-7977 *in vitro*. **(G-H)** RMC-7977-resistant cell lines derived from *in vivo* resistant PDX-4872 melanoma (C3M2, **G**) or RMC_7977^R^ cell line derived from parental PDX-4872 cell line under drug pressure *in vitro* (clone 6, **H**) at indicated concentrations. All graphs are representative of n=3 independent experiments. Error bars represent standard deviation (SD) of triplicate technical replicates.

Similarly, RMC-7997-sensitive or resistant NRAS^Q61R^/INK4A-ARF^Null^ melanoma cell lines were isolated from the *TNC* GEM model **(Fig. 4C-F)** leading to the generation of four cell lines: TNC-529 (from a vehicle treated mouse) and; TNC-409, TNC-527 and TNC-533 (from RMC-7977^R^ melanomas). Incucyte analysis of melanoma cell proliferation confirmed the sensitivity of TNC-529 cells, and the resistance (up to 1μM) of TNC-409, –527 and –533 cells. Sensitivity analysis to MEK1/2 or ERK1/2 inhibition, using trametinib or ulixertinib respectively, was performed to determine whether RMC-7977^R^ cells remain dependent on MAPK pathway signaling. Notably, the three RMC-7977^R^ lines exhibited cross-resistance to trametinib (100nM) or ulixertinib (1μM), whereas TNC-529 cells were sensitive to all three agents (Figs. 4C–F), suggesting that MAPK pathway reactivation at or downstream of MEK1/2 might be the driver of acquired resistance rather than bypass through parallel pathways.

Finally, we generated RMC-7977 sensitive or resistant human PDX-4872 melanoma cell lines from vehicle or compound-treated mice **(Fig. 4G)**. Proliferation of untreated PDX-4872 cells was inhibited by ≥10nM RMC-7977 **(Fig. 1F)**. By contrast, cells derived from resistant melanomas proliferated in ≤500nM RMC-7977 **(Fig. 4G).** It is interesting to note that sorting of single untreated PDX-4872 cells into the wells of a 96 well dish followed by treatment with 100nM RMC-7977 led to the isolation of nine resistant cell clones, five of which (4, 5, 6, 8 & 9) were expanded for further analysis, suggesting that the frequency of pre-existing RMC-7977^R^ cells in parental PDX-4872 cells is high. Indeed, some of these cells (e.g., clones 4 & 6) proliferated in 500nM RMC-7977 **(Fig. 4H)**. However, immunoblot analysis of these five clones revealed, in most cases, striking differences in the baseline levels of pERK1/2 and pAKT1-3 compared to the parental (P) cells (Suppl. Fig. S6A). While the significance of the basal phospho-protein levels is not entirely certain at this time, this may reflect a diversity of underlying molecular mechanisms driving RMC-7977 resistance in the parental tumor cell population.

### Mutational silencing of CYPA expression is a mechanism of RMC-7977 resistance

To identify mechanisms of resistance, we initially analyzed RMC-7977^R^ O.1 or O.10 cells: O.1^RM6^ and O.10^RM1^, respectively. Initial immunoblot analysis indicated that CYPA expression was silenced in these cells compared to parental cells **(Fig. 5A-B)**. Furthermore, immunohistochemical analysis of sections of RMC-7977^R^ tumors harvested at experimental endpoint **(Fig. 2A, C)** indicated that selection for CYPA silencing occurred in tumor-bearing mice **(Fig. 5C)**. Hence, to test if restoration of CYPA expression would restore sensitivity to RMC-7977, we ectopically expressed human CYPA in CYPA-deficient O.1^RM6^ cells, which was confirmed by immunoblotting at levels comparable to parental OSUMMER.1 cells **(Fig. 5D)**. Incucyte analysis revealed that ectopic expression of CYPA re-sensitized O.1^RM6^ cells to RMC-7977 **(Fig. 5E)**.

**FIGURE 5:**
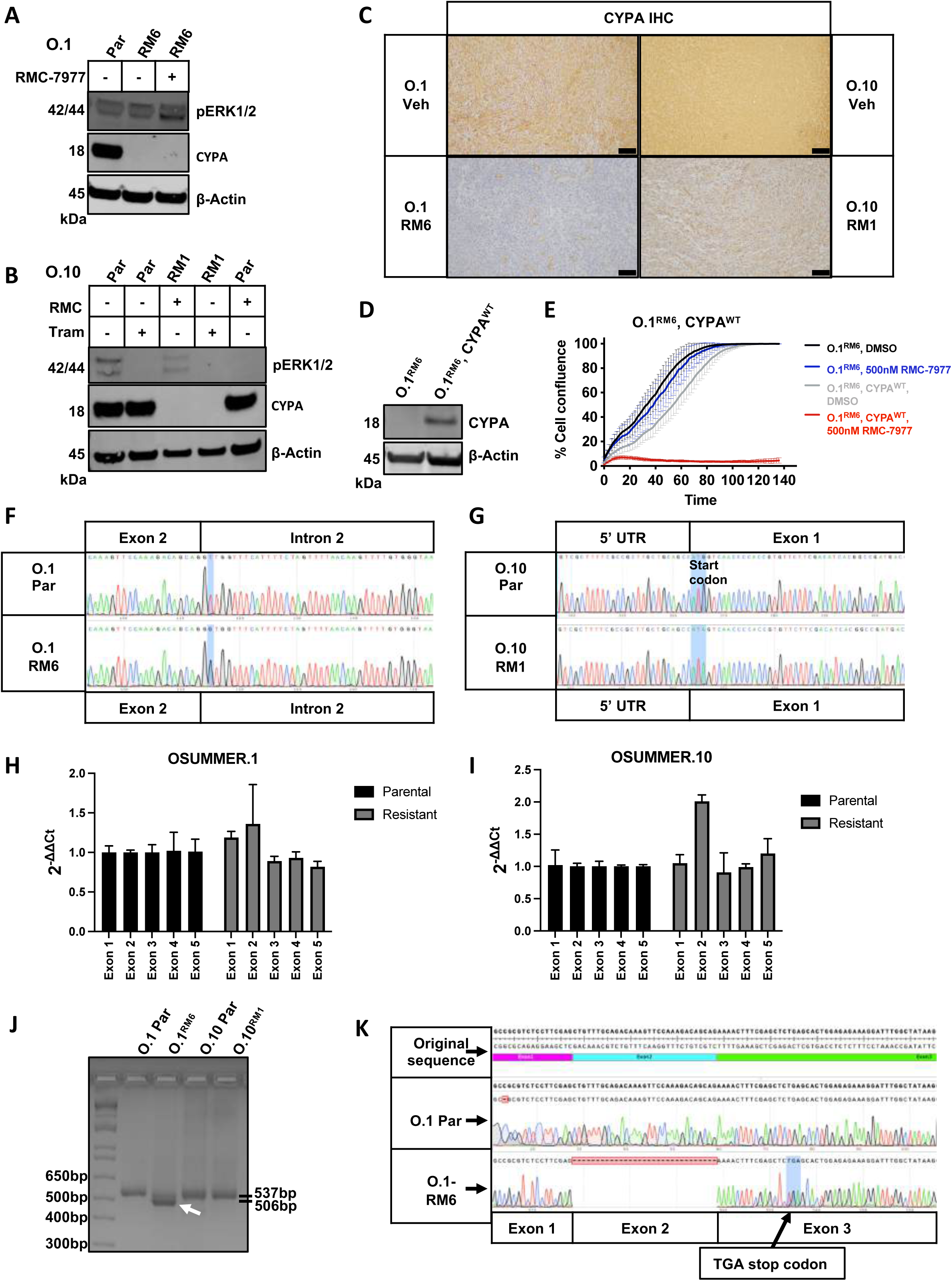
Mutational silencing of CYPA expression is a mechanism of RMC-7977 resistance. **(A-B)** Immunoblots of lysates from OSUMMER.1 (NRAS^Q61R^)-derived RMC_7977^R^ cell line O.1^RM6^ **(A)** or OSUMMER.10 (NRAS^G13R^)-derived RMC_7977^R^ cell line O.10^RM1^ **(B)** versus parental (Par) cell lysates probed for the presence or absence of total CYPA protein or phospho-protein ERK1/2 and β-actin. **(C)** Immunohistochemistry (IHC) for CYPA on representative sections of tumors from mice (O.1-Veh vs. O.1^RM6^; or O.10-Veh vs. O.10^RM1^; n=3 each) treated with vehicle (O.1, 12 days; O.10, 10 days) or RMC-7977 until resistance (O.1, 24 days; O.10, 32 days; dosing experiments depicted in Fig. 2A, C). Scale bar in lower right corner represents 100uM in all images. **(D)** Immunoblot for CYPA protein upon ectopic expression of human wildtype Cyclophilin A (CYPA^WT^) in O.1^RM6^. **(E)** Cell confluency assay of O.1^RM6^ cells (O.1^RM6^) and O.1^RM6^ cells ectopically expressing CYPA^WT^ (O.1^RM6^-CYPA^WT^) under RMC-7977 treatment at indicated concentration using an Incucyte Live-Cell Analysis System. Graph is representative of n=3 independent experiments. Error bars represent standard deviation (SD) of triplicate technical replicates. **(F)** Sanger sequencing of Parental OSUMMER.1 or RMC_7977^R^ O.1^RM6^ cells. Blue highlighted base is mutated in O.1^RM6^ (T>G). **(G)** Sanger sequencing of Parental OSUMMER.10 or RMC_7977^R^ O.10^RM1^ cells. Blue highlighted start codon (ATG) is mutated in O.10^RM1^ to AGA (G>A). **(H, I**) Quantitative PCR (qPCR) on genomic DNA on each *Ppia* exon from O.1 **(H)** or O.10 **(I)** parental vs. O.1^RM6^ (**H)** or O.10^RM1^ **(I)** RMC_7977^R^ cell lines as indicated. **(J)** PCR amplification of exon 2 from cDNA of parental OSUMMER.1 or 10 vs. O.1^RM6^ or O.10^RM1^. **(K)** Sanger sequencing of gel extracted bands from exon 2 amplified PCR products of parental O.1 vs. O.1^RM6^. Novel TGA stop codon in exon 3 due to frameshift is highlighted in blue.

Targeted sequencing of the five coding exon *Ppia* gene (encoding CYPA) in parental vs O.1^RM6^ cells revealed a single point mutation (G<u>T</u>>G<u>G)</u> in the intron 2 splice donor **(Fig. 5F)**. This point mutation appeared to be either homozygous or hemizygous, indicated by a single, non-overlapping sequencing peak at position c.100+2(T>G) **(Fig. 5F)**. Furthermore, DNA sequence analysis of the *Ppia* locus in O.10^RM1^ revealed a point mutation at the translation initiation codon in exon 1 of *Ppia* mRNA (AT<u>G</u>>AT<u>A</u>) at position c.3(G>A), which also appeared to be homozygous or hemizygous **(Fig. 5G).** Although there is an upstream and in-frame AUG codon (position c.-412-414(ATG)) within the 5’-UTR of mouse *Ppia* that would, in principle, promote translation of a 138 amino acid N-terminally extended form of CYPA, we have not detected expression of such an elongated form of CYPA using two different CYPA antibodies that recognize residues at the C– or N-terminus of CYPA. Hence, it seems likely that the splice donor or translation initiation mutations in O.1^RM6^ and O.10^RM1^ cells respectively are the reason why CYPA is silenced and the cells are therefore RMC-7977 resistant.

To determine the zygosity of the *Ppia* mutations, we performed quantitative PCR (qPCR) using genomic DNA (gDNA) from parental versus O.1^RM6^ or O.10^RM1^ cells **(Fig. 5 H-I)**. Targeted amplification of *Ppia* exons 1-5 revealed similar copy number between parental and resistant clones **(Fig. 5 H-I)**. Importantly, amplification of an X-chromosome-linked gene (GATA1) demonstrated that our qPCR technique can accurately detect the difference between one or two alleles of a gene **(Suppl. Fig. S6B-C).** Moreover, karyotype analysis of O.1^RM6^ and O.10^RM1^ cells revealed that they retained two copies of chromosome 11, the location of mouse *Ppia* **(Suppl. Fig. S6 D, E).** These data suggest that the O.1^RM6^ and O.10^RM1^ retain two copies of the *Ppia* gene and that the inactivating mutations that silence CYPA expression are identical in both copies.

To determine how the *Ppia* intron 2 splice donor mutation would affect the expression of *Ppia* mRNA, we designed primers to amplify exons 1, 2 or 3 from cDNA. We hypothesized that intron 2 sequences might be retained in the mature mRNA sequence, resulting in a larger, incompletely spliced mRNA. Contrary to our hypothesis, PCR analysis yielded a shorter band in O.1^RM6^ mRNA in which DNA sequence analysis revealed that *Ppia* mRNA had been aberrantly spliced from the intron 1 splice donor to the intron 3 splice acceptor thereby entirely emitting the 31bp exon 2 leading to expression of *Ppia^ΔEx2^* mRNA **(Figs. 5 J-K)**. Translation of the predicted *Ppia^ΔEx2^*mRNA would be terminated by a frameshift that inserts a premature UGA stop codon at position c.115-118 of exon 3, allowing for translation of a polypeptide of 28 amino acids, which was not detectable by immunoblotting using antisera directed against the N-terminus of CYPA.

### CYPA-deficient melanoma cells are tumorigenic in immunocompetent mice

CYPA is a cis-trans prolyl-isomerase, which has been suggested to be essential for cell proliferation through effects on protein folding, trafficking and/or cell cycle regulation (11,12,32,33). Although CYPA-deficient OSUMMER cells grow rapidly in culture, we tested whether they are capable of forming tumors in mice. To that end we injected 10^6^ of the following CYPA-deficient cells into *GH* mice: O.1^RM6^; O.10^RM1^; and O.1^CYPA-CRISPR^, the last of which was generated by silencing CYPA expression using CRISPR/CAS9 gene editing **(Suppl. Fig. S7 A-D)**. All three cell lines rapidly formed tumors comparable to that of CYPA-proficient parental O.1 or O.10 cells (∼1-2 weeks). Tumor lysates were analyzed by immunoblotting using antibodies that recognize either the N– or C-terminus of CYPA **(Fig. 6A)** thereby revealing that CYPA-deficient OSUMMER cell-derived tumors remained CYPA-negative. The small amount of CYPA detected in these blots likely originates from CYPA proficient normal mouse stroma cells within the tumor microenvironment **(Fig. 6A).** This observation was further confirmed by immunohistochemistry where CYPA expression was substantially absent from O.1^RM6^ or O.10^RM1^ tumor cells compared to cells in vehicle-treated O.1 or O.10 parental tumors **(Fig. 6B).**

**FIGURE 6:**
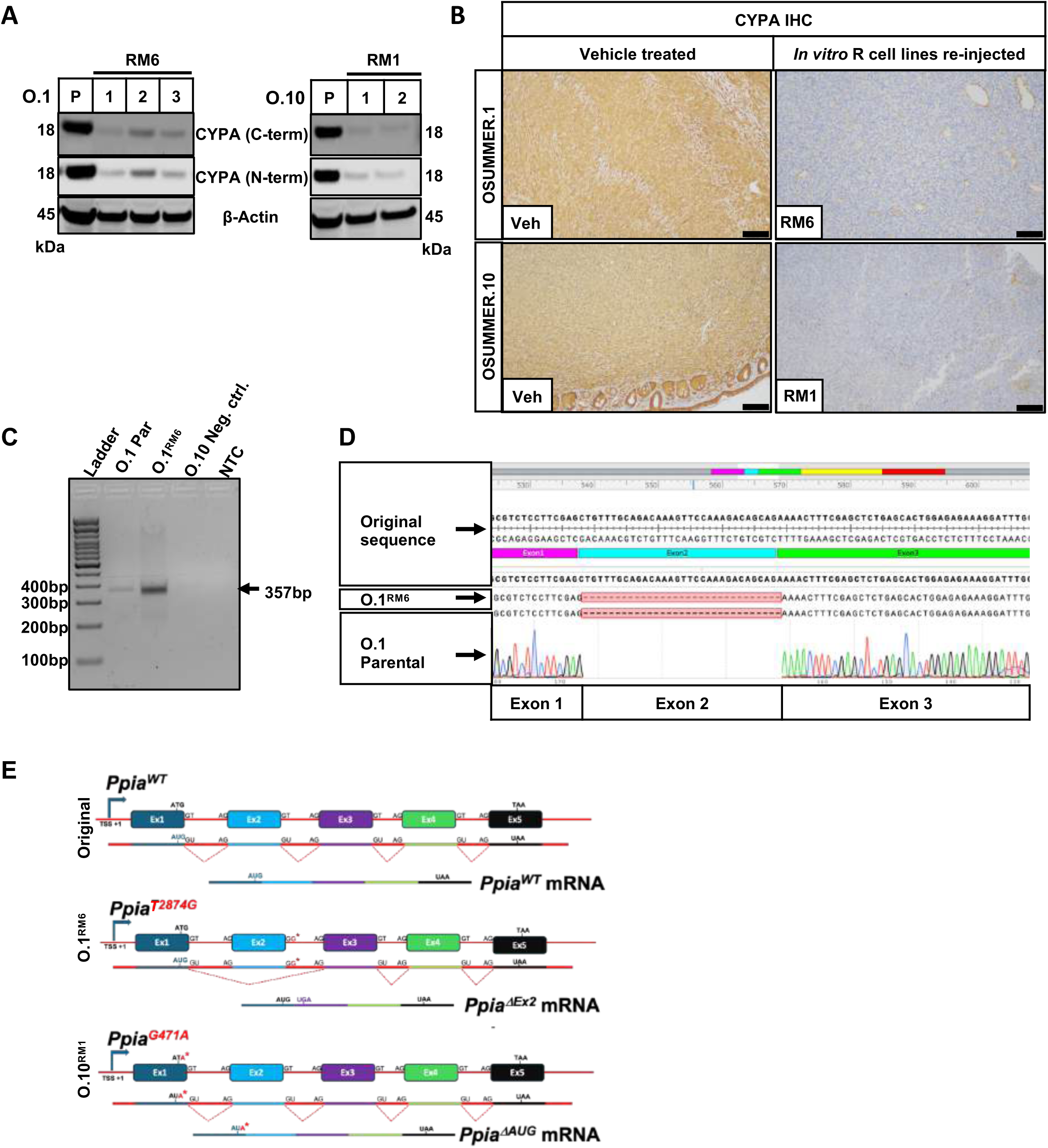
CYPA-deficient melanoma cells are tumorigenic in immunocompetent mice. **(A)** Immunoblot for total CYPA protein (antibodies recognizing C-terminus vs. N-terminus) or β-actin, on lysates from tumors forming in mice re-injected with RMC_7977^R^ O.1^RM6^ or O.10^RM1^ cells. **(B)** IHC for CYPA on representative sections of tumors from mice re-injected with RMC_7977^R^ O.1^RM6^ or O.10^RM1^ cells. Scale bar in lower right corner represents 100uM in all images. **(C)** PCR amplification of novel exon 1-3 junction in parental O.1 (O.1 Par) or O.1^RM6^ cDNA. O.10 and a non-template control (NTC) served as negative controls. **(D)** Sanger sequencing of gel extracted bands from exon 1-3 junction PCR products at 357bp. **(E)** Schematic of genetic alterations in the *Ppia* gene of O.1-RM6 (*Ppia^T2874G^*) or O.10-RM1 (*Ppia^G471A^*) compared to the original sequence (*Ppia^WT^*), resulting in alternative mRNA transcripts *Ppia^ΔEx2^* or *Ppia^ΔAUG^* compared to *Ppia^WT^* mRNA, respectively.

To determine if cells expressing the aberrantly spliced *Ppia^ΔEx2^* mRNA pre-exist in parental OSUMMER.1 cells prior to RMC-7977 treatment, we designed a PCR strategy that would detect the novel exon1⇔exon3 junction in the *Ppia^ΔEx2^* mRNA expressed in O.1^RM6^ cells **(Fig. 6C).** As a negative control we used mRNA from parental

O.10 cells, which should not contain the *Ppia^ΔEx2^* mRNA. By this strategy we could readily detect *Ppia^ΔEx2^* mRNA in both the RMC-7977^R^ O.1^RM6^ cells, but also in untreated parental O.1 cells. As anticipated, *Ppia^ΔEx2^* mRNA was undetectable in parental O.10 cells **(Fig. 6C)**. Targeted sequencing of the PCR product derived from the parental O.1 cells (lane 1) confirmed the absence of the 31bp exon 2 sequence, exactly as documented in O.1^RM6^ cells (lane 2) (**Fig. 6D)**. These data confirm that cells expressing the aberrantly spliced *Ppia^ΔEx2^* mRNA pre-exist in the parental O.1 population and are not generated after or in response to RMC-7977 exposure. Overall, these data indicate that bi-allelic silencing of *Ppia/CYPA* expression is a mechanism of RMC-7977 resistance **(Fig. 6E)**.

### *Map2k1* gain-of-function mutations are an alternate mechanism of RMC-7977 resistance

While some RMC-7977 resistant cells were CYPA deficient, others retained CYPA expression (Suppl. Fig. S8A-B). To identify additional mechanisms of resistance, we subjected CYPA expressing, RMC-7977^R^ O.1, O.10 or PDX-4872 cell lines to whole exome sequencing (WES). Of the six analyzed OSUMMER-derived cell lines, two carried *Ppia* missense mutations, resulting in expression of CYPA^M136R^ or CYPA^C62W^, with an additional cell line carrying a *Ppia* frameshift mutation resulting in CYPA^H126X^ **(Fig. 7A; Suppl. Table S4, S5).** The other four cell lines each carried well-known hotspot mutations in *Map2k1* (encoding the RAF-regulated, dual-specificity protein kinase, MEK1), specifically MEK1^K57T^ (O.1^RM5^), MEK1^K57N^ (O.1^RM6-B^), MEK1^Q56P^ (O.10^RM3^) or MEK1^C121S^ (O.10^RM4^, **Fig. 7A**, (34,35)). It should be noted that O.1^RM6-B^ cells were isolated from the same tumor as the CYPA-deficient O.1^RM6^ cell line. However, unlike O.1^RM6^ cells that are resistant to 1μM RMC-7977, O.1^RM6-B^ cells are resistant to a maximum of ∼100nM RMC-7977 indicating that these two cell lines stemmed from two different clones harboring different mechanisms of RMC-7977 resistance within the same tumor.

**FIGURE 7:**
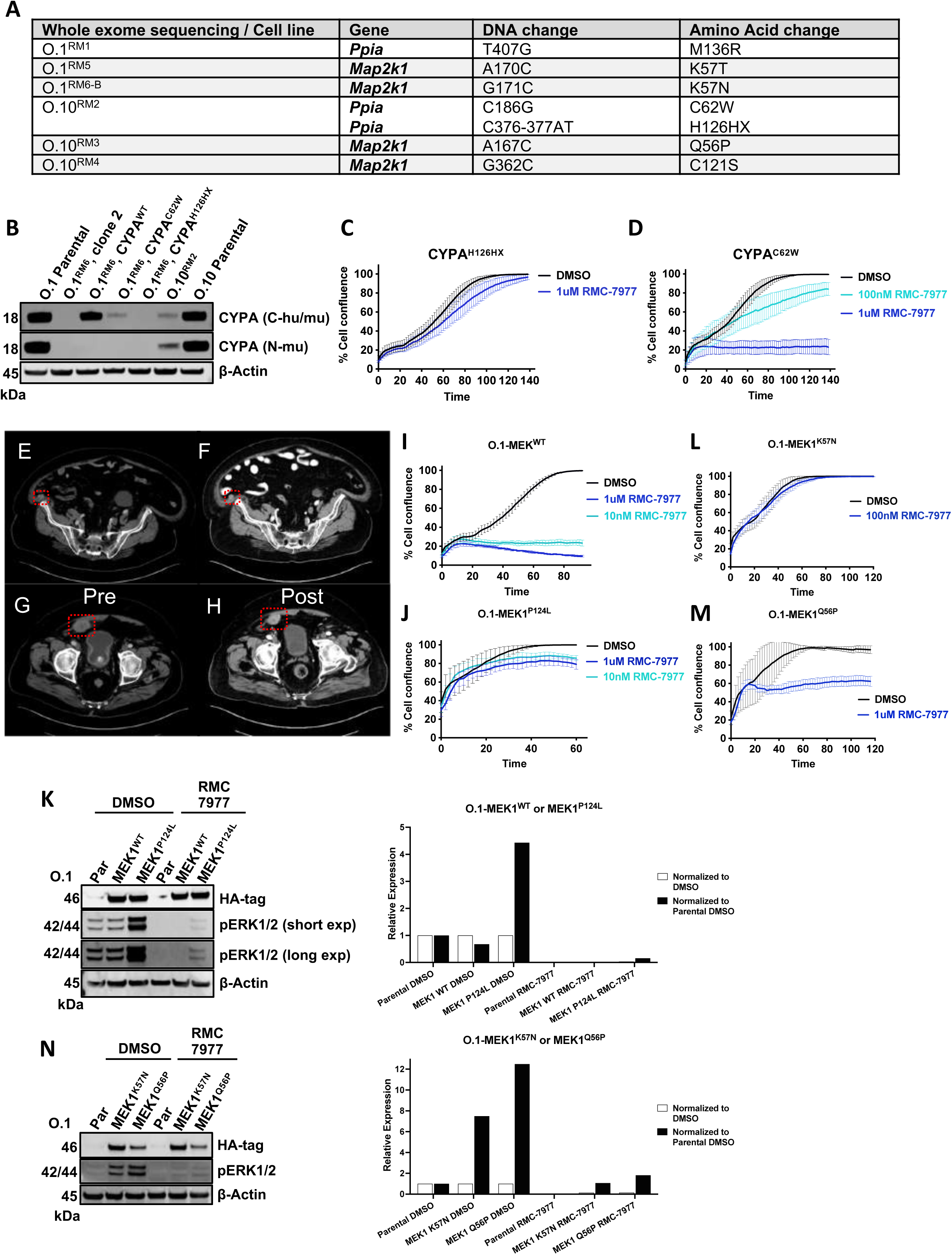
M*a*p2k1 gain-of-function mutations mediate RMC-7977 resistance. **(A)** Whole exome sequencing (WES) on three RMC_7977^R^ cell lines derived from OSUMMER.1 (NRAS^Q61R^; RM1, RM5, or RM6-B) or OSUMMER.10 (NRAS^G13R^; RM2, RM3, or RM4) identifying *Ppia* or *Map2k1* mutations and their corresponding amino acid changes as indicated. **(B-D, I-N)** Re-expression of *Ppia* mutations identified by WES in CYPA-deficient cell line O.1-RM6. **(B)** Immunoblot for total CYPA protein on lysates from parental OSUMMER.1, O.1^RM6^ clone 2, O.1^RM6^-CYPA^WT^, O.1^RM6^-CYPA^H126HX^ or CYPA^C62W^. O.10^RM2^ is the original cell line that expressed both CYPA^H126HX^ and CYPA^C62W^ that were identified by WES. The C-terminus CYPA antibody recognizes mouse and human (C-hu/mu), while the N-terminus CYPA antibody is mouse-specific (N-mu). The CYPA expression vector *pScalps-huCypA-WT* is a human construct. **(C, D)** Cell confluency assay of O.1-RM6 CYPA^H126HX^ **(C)** or CYPA^C62W^ **(D)** treated with RMC-7977 as indicated and analyzed by Incucyte Live-Cell Analysis. Graphs are representative of n=3 independent experiments. Error bars represent standard deviation (SD) of triplicate technical replicates. **(E-H)** Computed tomography (CT) images of an NRAS^Q61K^-driven melanoma patient before (*pre,* **E, G)** and after 18 weeks on treatment with 300mg daraxonrasib, q.d. (*post*; **F, H)**. Red dotted boxes indicate location of the metastatic target lesions. Target lesion 1: L 22.8 mm x W 15.7 mm **(E)**, L13.9 mm x W 12.1 mm **(F)**. Target lesion 2: L 41.6 mm x W 28.3 mm **(G)**, L 30.7 mm x W 26.0 mm **(H)**. **(I, J, L, M)** Cell confluency assays of OSUMMER.1 ectopically expressing MEK1^WT^ **(I),** MEK1^P124L^ **(J),** MEK1^K57N^ **(L),** or MEK1^Q56P^ **(M)** under concentrations of RMC-7977 as indicated and analyzed by Incucyte Live-Cell Analysis. Graphs are representative of n=3 independent experiments. Error bars represent standard deviation (SD) of triplicate technical replicates. **(K, N)** Immunoblots for HA-MEK1 fusion protein, pERK1/2 (short or long exposure; exp), or β-actin on lysates from parental O.1 (Par), or O.1 ectopically expressing MEK1^WT^ or MEK1^P124L^ **(K),** or MEK1^K57N^ or MEK1^Q56P^ **(N)** in the absence or presence of 100nM RMC-7977 as indicated.

To date, WES analysis of four RMC-7977 resistant PDX-4872-derived cell lines failed to reveal any genetically hard-wired mechanisms of resistance **(Suppl. Table S6).** However, we have detected differences in DNA sequence between parental and RMC-7977^R^ PDX-4872 cells in genes involved in chromatin organization, transcriptional regulation, protein folding, protein ubiquitination, energy metabolism, autophagy, calcium signaling, or antigen processing and presentation, reflecting broader cellular re-arrangements that could have contributed to the development of resistance in the RMC-7977^R^ PDX-4872 cells **(Suppl. Table S6).** Notably, no obvious PI3’-kinase>AKT pathway-related genes were mutationally altered in any of the RMC-7977^R^ cells that were sequenced. However, we tested the hypothesis that elevated flux through the PI3’-kinase pathway might confer RMC-7977 resistance by analysis of O.1 cells engineered to express an activated form of AKT1 **(Suppl. Fig. S9A, B)**, or in O.1 cells in which PTEN was silenced by CRISPR/CAS9 gene editing **(Suppl. Fig. S9 C-I)**, or a mutationally-activated RAC1 (RAC1^P29S^, **Suppl. Fig. S9 J-M**) (36–39). All of these cells remained equivalently sensitive to the anti-proliferative effects of RMC-7977 **(Suppl. Fig. S9 B, D-F, L-M).**

To confirm that the point mutations detected in *Ppia* that result in expression of CYPA^H126HX^, CYPA^C62W^, or CYPA^M136R^ contribute to RMC-7977 resistance, we tested whether lentivirus-mediated expression of these CYPA variants could restore RMC-7977 sensitivity in CYPA deficient O.1^RM6^ cells. Expression of the CYPA variants was assessed by immunoblotting using two CYPA antibodies recognizing either the C– or N-terminus as described above. The C-terminally directed antibody detects both human and mouse CYPA (C-hu/mu) whereas the N-terminally directed antibody only detects murine CYPA (N-mu) **(Fig. 6A)**. Although we could detect expression of the normal CYPA as well as CYPA^C62W^ **(Fig. 7B)** and CYPA^M136R^ **(Suppl. Fig. S10A)** we could not detect CYPA^H126HX^ (frameshift) **(Fig. 7B).** Moreover, there was no change in sensitivity of O.1^RM6^ cells engineered to express CYPA^H126HX^ encoding lentivirus (growth inhibition, GI_50_>1uM), and substantial resistance to RMC-7977 was observed in O.1^RM6^ cells engineered to express CYPA^C62W^ (GI_50_ = 207nM), further confirming that the CYPA^H126HX^ and CYPA^C62W^ variants are compromised in their ability to form a functionally inhibitory RMC-7977⇔CYPA⇔RAS⇔GTP inhibited complex **(Fig. 7C, D; Suppl. Fig. S10A).** Finally, expression of CYPA^M136R^ substantially sensitized O.1-RM6 cells to RMC-7977 **(Suppl. Fig. S10B),** suggesting that this variant is not the primary driver of resistance in the cells in which it was detected.

### Case study 1: A patient with NRAS^Q61K^-driven metastatic melanoma responded to daraxonrasib as evidenced by regression of at least two metastatic lesions

As preclinical research was ongoing, a patient (#1) with metastatic (lungs, peritoneum and in-transit limb lesions) NRAS^Q61K^-driven melanoma was enrolled into an *NRAS*-mutated melanoma-specific expansion cohort of the Phase 1 clinical trial of daraxonrasib (NCT05379985). The patient, an 82 year old male, had progressed on prior immunotherapy (pembrolizumab and TVEC) and received daraxonrasib treatment at 300mg daily with concurrent doxycycline prophylaxis for skin rash. In cycle 1, Patient #1 tolerated daraxonrasib with grade 1 adverse events (nausea, diarrhea). After cycle 2, treatment was reduced to 200mg daraxonrasib following grade 2 adverse effects (stomatitis, skin rash). After cycle 10, a second dose reduction to 160mg was required due to worsening skin rash. A re-staging CT scan after cycle 6 showed a partial response (PR; –39% per RECIST; **Fig. 7E-F** and **G-H**) and restaging after cycle 14 continued to show a PR (–44%). Following cycle 15, skin disease progression was noted but deemed not to be clinically significant, allowing treatment to continue per protocol. The response to daraxonrasib treatment observed in Patient #1 highlights the potential impact of daraxonrasib-mediated direct inhibition of RAS[GTP] signaling in *NRAS*-driven melanoma even with dose modifications in an elderly and heavily pre-treated patient.

### Case study 2: A pre-existing MEK1^P124L^ alteration is the likely cause of resistance to daraxonrasib in a patient with *NRAS*-mutated melanoma

A second patient (#2), a 43-year old female with metastatic (lung and chest wall) NRAS^Q61K^-driven melanoma, who had progressed after prior cycles of immunotherapy (nivolumab plus relatlimab), was enrolled into the same trial as Patient #1. Patient #2 was treated with daraxonrasib at 300mg daily in 21-day cycles, exhibiting grade 1 or 2 adverse events (stomatitis, rash, intermittent diarrhea) on treatment. A computed tomography (CT) scan prior to cycle 3 showed stable disease (+3% per RECIST) as best response to therapy. However, a re-staging CT scan prior to cycle 7 showed disease progression in the lungs and chest wall (+26% per RECIST) at which point treatment was discontinued. Analysis of baseline NGS from the tumor indicated that, prior to daraxonrasib treatment, in addition to the *NRAS^C181A^*(VAF: ∼48%) encoding NRAS^Q61K^ plus a *CDKN2A* loss-of-function alteration, the patient also had a *MAP2K1^C371T^* mutation (VAF: ∼38%) encoding MEK1^P124L^. Moreover, this co-mutation of *NRAS* (VAF51.1%) and *MAP2K1* (VAF=34.4%) was detected in DNA isolated from the patient’s diagnostic specimen prior to any therapy. MEK1^P124L^ is a noted gain-of-function alteration that is: 1. Detected at low frequency in certain cancers, sometimes in combination with *NRAS*; 2. A driver of Cardio-Facio-Cutaneous (CFC) syndrome and; 3. A mediator of resistance of BRAF^V600E^-driven melanomas to either RAF or MEK inhibitors (17–19).

To test if MEK1^P124L^ can confer RMC-7977 resistance, we ectopically expressed either MEK1^WT^ or MEK1^P124L^ in O.1 **(Fig 7I-K)** or O.10 cells **(Suppl. Fig. S10C, D, E).** While expression of MEK1^WT^ had little or no effect on the sensitivity of O.1 cells to RMC-7977 (GI_50_: 5.9nM), expression of MEK1^P124L^ rendered O.1 or O.10 cells substantially RMC-7977 resistant (O.1 GI_50_: 664nM; O.10 GI_50_: 2,495nM) **(Fig. 7I-J; Suppl. Fig. S10C-D).** Immunoblot analysis of parental versus MEK1^P124L^ expressing O.1 or O.10 cells indicated that mutationally-activated MEK1^P124L^ sustained elevated pERK1/2, albeit at diminished levels, in the presence of RMC-7977 **(Fig. 7K; Suppl. Fig. S10G)**, consistent with MEK1^P124L^ remaining at least partly dependent on upstream RAS-GTP signaling (34). These data strongly suggest that the resistance to daraxonrasib observed in Patient #2 was (at least in part) driven by MEK1^P124L^ co-expression at baseline in the majority of the NRAS^Q61K^-driven melanoma cells. Moreover, we tested whether expression of MEK1^K57N^ or MEK1^Q56P^, two variants that were found in *Map2k1* by WES **(Fig 7A),** would result in resistance to RMC-7977. Both variants rendered O.1 or O.10 cells substantially resistant to RMC-7977, i.e. 100nM (O.1 MEK1^K57N^; GI_50_ 125.1nM **(Fig. 7L; Suppl. Fig. S10E)** or 1μM (O.1 MEK1^Q56P^; GI_50_ 898.4nM) **(Fig. 7M; Suppl. Fig. S10F)**. Similar to MEK1^P124L^, immunoblot analysis of parental versus MEK1^K57N^ or MEK1^Q56P^ expressing O.1 or O.10 cells indicated that mutationally-activated MEK1^K57N^ or MEK1^Q56P^ sustained elevated pERK1/2, albeit at diminished levels, in the presence of RMC-7977 **(Fig. 7N; Suppl. Fig. S10H).** These findings, together with recent data identifying *MAP2K1* mutations resulting in MEK1-K57N, –K57T, or –Q56P in daraxonrasib resistant patient samples (40), strongly support *MAP2K1* mutation as a recurrent, clinically relevant resistance mechanism to RAS(ON) inhibition.

## DISCUSSION

Melanoma therapy has been revolutionized by FDA-approved pathway-targeted or immunotherapies, as well as by T-cell-based or oncolytic virus therapies (41–44). However, ∼80% of patients with NRAS-driven melanoma, particularly those ineligible for, or resistant to immune checkpoint inhibitors, lack effective second-line treatment options (45,46). Moreover, efforts to target RAS pathway signaling downstream of NRAS, most notably the RAF>MEK>ERK and/or the PI3’-kinase>AKT pathways have been limited by lack of efficacy, toxicity or both (5–7). Recently, the discovery of direct RAS(ON) inhibitors such as daraxonrasib offers a new avenue for potent anti-melanoma therapeutic options (9,10). Indeed, our preclinical models of NRAS-driven melanoma initially display exquisitely specific and selective RMC-7977 sensitivity. Further to this, daraxonrasib has been evaluated in a first-in-human clinical trial with expansions into different cohorts including patients with NRAS-driven melanoma (NCT05379985). Initial results in this Phase 1 trial demonstrated the efficacy and safety of daraxonrasib at doses up to 300mg/day with most adverse events limited to low grade toxicities including skin rash (90%), stomatitis (30%) and diarrhea (48%) (47). The daily dose of 300mg was selected for further evaluation in ongoing global randomized Phase 3 clinical trials in patients with metastatic *RAS*-mutated pancreatic cancer or non-small cell lung cancer (RASolute 302: NCT06625320; RASolve 301: NCT06881784).

However, as with many single-agent pathway-targeted cancer therapies, primary or acquired resistance may limit the depth and durability of patient responses (48). Indeed, mechanistic insights from our preclinical models prompt two conclusions: 1. CYPA is essential for this class of RAS(ON) inhibitor’s mechanism of action as shown previously (9,10,14), and; 2. The primacy of RAF>MEK>ERK signaling for the maintenance of NRAS-driven melanoma (Suppl. Table S7). Indeed, we show here that hot-spot gain-of-function mutations in *BRAF* or *Map2k1* (encoding MEK1) are a mechanism of primary or acquired resistance to RMC-7977 respectively. Finally, we show that a gain-of-function *MAP2K1* mutation encoding MEK^P124L^ is a likely candidate resistance mechanism observed in Patient #2 with NRAS^Q61K^-driven melanoma. Although there is evidence suggesting that PI3’-kinase>AKT signaling can confer resistance to inhibitors of BRAF^V600E^ signaling in melanoma (49–51), we see variable modulation of this pathway in response to RMC-7977, and no evidence that deliberate activation of this pathway renders NRAS-driven melanoma cells RMC-7977 resistant. It should be noted that elevated c-MYC expression as a mechanism of resistance to RMC-7977 or to the combination of trametinib plus hydroxychloroquine in KRAS-driven pancreatic cancer as previously described (15,52), was not observed in our NRAS-driven melanoma models. Despite the essentiality of RAF>MEK>ERK signaling in the maintenance of NRAS-driven melanoma, FDA-approved MEK1/2 inhibitors (e.g., trametinib or binimetinib) have shown limited clinical efficacy against NRAS-driven melanoma, often attributed to concurrent PI3’-kinase>AKT pathway activity (53–55). However, our analyses suggest that RMC-7977 does not have immediate and direct inhibitory effects on PI3’-kinase>AKT signaling in this context, and that mutational activation of this pathway does not confer RMC-7977 resistance in our models. These data suggest that there may be additional signaling pathways that obviate sensitivity of NRAS-driven melanomas to MEK1/2 inhibitors. A potential candidate is YAP/TAZ>TEAD signaling, known to confer resistance to RAS or BRAF inhibitors in other contexts (15,56,57), or the recently described effects of RAS oncoproteins on nuclear export via RAN GTPases (58). Additional analyses will be required to reveal differences in the biochemical and biological responses of NRAS-driven melanoma to either single-agent RAS or MEK1/2 inhibitors.

There remain challenges in the development of predictive biomarkers of response or resistance to daraxonrasib in melanoma beyond *NRAS* mutations. Notably, CYPA silencing requires bi-allelic silencing of *PPIA*. CRISPR screens in DepMap (https://depmap.org/portal) across many cancer cell lines indicate that *PPIA* falls into the moderate-to-essential gene category (<u>C</u>omputational <u>E</u>stimation of <u>R</u>elative <u>E</u>ssentiality <u>S</u>cores (CERES) score is ∼0.5, compared to –1 for essential genes). However, the essentiality of *PPIA* may vary depending on the cell type, growth conditions and the expression of one or more of the other 17 human *PPI* genes encoding cis-trans prolyl-isomerases (11,59). Certainly, in our experiments, *Ppia* was dispensable for O.1 or O.10 cell proliferation or tumorigenesis, consistent with previous observations (9,32). It is interesting to note that the inactivating splice donor or initiator ATG codon mutations in O.1^RM6^ or O.10^RM1^ cells respectively were not signature UV alterations (C>T, CC>TT), and appeared to be homozygous (i.e., identical mutations in both copies of the same gene), which might be considered a low frequency spontaneous event. These data suggest that an initial deleterious mutation in one copy of *Ppia* was then subject to copy-neutral loss of heterozygosity either by gene conversion or the process of uniparental disomy (60). It will therefore be interesting to determine the frequency and mechanism(s) of *PPIA*/CYPA silencing in patients as a mechanism of daraxonrasib resistance. Finally, silencing of *PPIA* in daraxonrasib resistant cells may present an opportunity for a synthetic lethality strategy to target cells lacking CYPA expression. Data from previously published preclinical models of pancreatic cancer and data presented here suggest that combination therapy will likely be superior to RAS inhibition alone (20,22). Preclinical models, such as the OSUMMER *NRAS*-mutated melanoma cells in immunocompetent mice will allow the evaluation of the preclinical anti-tumor activity of combination therapies such as RAS(ON) multi-selective plus immune checkpoint inhibitors with obvious translational potential (61).

## MATERIALS & METHODS

### Cell Culture

Mouse NRAS-driven melanoma cell lines: OSUMMER.1; OSUMMER.8; OSUMMER.10 and; OSUMMER.11 **(Suppl. Table S2)** were obtained from Millipore and maintained in DMEM (Invitrogen) supplemented with 10%(v/v) fetal bovine serum (FBS) and 1% penicillin/streptomycin (23). Human NRAS-driven melanoma PDX-derived cell lines: MTG2; Mel9 and; PDX-4872 **(Suppl. Table S2)** were generated in-house and maintained in Mel-2 medium **(Suppl. Table S8)** supplemented with 2%(v/v) FBS and 1% antibiotics. STR analysis was performed on early passage human melanoma cell lines. For karyotyping, early-passage RMC-7977 resistant OSUMMER lines were submitted to Cell Line Genetics for G-banded karyotyping. Ten metaphase spreads from each cell line were karyotyped, and chromosomal aberrations were noted in the manuscript. To generate cell cultures made from primary tumors, tissue chunks were dissociated using the Miltenyi Mouse Tumor dissociation kit (#130-096-730) as per manufacturer guidelines. Cultures were monitored every 3 months for mycoplasma contamination using a PCR-based assay. All experiments were performed using cell lines kept in culture for no more than 10 passages.

### Cell Confluency, cytotoxicity, or PRISM Assays

Cell confluency or cytotoxicity was assessed by seeding 5-8 x 10^3^ cells/well in 96-well plates. Experiments were performed on two to three separate occasions in triplicate wells per experimental condition. Pharmacological agents were added 24 hours after cell seeding (treatment naive cell lines) or at seeding (resistant cell lines). Cells were cultured in the absence or presence of pharmacological agents for 3-5 days or until full confluency. Inhibitor concentrations used for each treatment are indicated in the figures or figure legends. For cytotoxicity assays, CytotoxRed (Sartorius # 4632) reagent was added 24 hours after seeding, at the time of compound administration. Confluence was assessed over time using an IncuCyte S3 instrument and IncuCyte Analysis Software (Essen BioScience) in two-hour intervals. For cytotoxicity assays, red object confluence (percent) was divided by the brightfield cell confluence (percent) per well, and then the ratio values were normalized to zero (0) hours per condition. For PRISM assay, RMC-7977 was screened against 931 PRISM DNA-barcoded cell lines established by the Broad Institute, as described in Holderfield et al. (9). The “RAS-PathwayOther” category is defined as cell lines with oncogenic or likely oncogenic mutations (according to OncoKB) in any of these genes: EGFR, ERBB2, ERBB3, ERBB4, MET, PDGFRA, FGFR1, FGFR2, FGFR3, FGFR4, KIT, IGF1R, RET, ROS1, ALK, FLT3, NTRK1, NTRK2, NTRK3, JAK2, CBL, ERRFI1, ABL1, SOS1, NF1, RASA1, PTPN11, KRAS (other than G12X), HRAS, NRAS, RIT1, ARAF, BRAF, RAF1, RAC1, MAPK1, MAP2K1, MAP2K2.

### Lentivirus Production

HEK293T cells were transfected with the relevant lentiviral vector(s) in combination with psPAX2 and pMD2.G lentiviral packaging plasmids. Viral supernatants were collected at 48 and 72 hours post transfection, filtered, and stored in aliquots at −80°C. Cells were transduced with 2-3 changes of lentiviral supernatant in the presence of 8μg/ml polybrene. Lentivirus infected cells were selected for 5 days with puromycin at 1-10μg/ml. Ectopic expression of lentiviral encoded proteins was confirmed by immunoblotting as described or by targeted sequencing.

### Mice and genotyping

All experimental procedures were approved by the University of Utah Institutional Animal Care and Use Committee (IACUC) and conducted in compliance with the facility’s animal welfare guidelines and protocols. Both male and female mice, 6-16 weeks of age, were used in this study. In the genetically engineered *Tyr::CreER^T2^; Nras^LSL-Q61R/LSL-Q61R^; Cdkn2a^lox/lox^* (*TNC*) mouse model, we generated two cohorts of mice with different tumorigenesis induction schedules: 1. Cre-mediated recombination and UVB-induced tumorigenesis was initiated in neonatal mice 1-2 days after birth (details in Suppl. **Fig. S4)** and 2. Cre-mediated recombination was initiated in adult mice at age 6-16 weeks. *TNC* mice and *Tyr::CreER^T2^; Braf^CA/+^; Cdkn2a^lox/lox^* (*TBC*) mice were maintained on a mixed C57Bl/6 and FVB/N background by random interbreeding. DNA from tail biopsies was used to genotype for the *Tyr::CreER^T2^* transgene or the various genetically modified *Braf^CA^*, *Cdkn2a^flox^* or *Nras^LSL-Q61R^*alleles as previously described **(Suppl. Table S9)** (27,28,62,63). Whilst the experiments were underway, we noted that some *TNC* mice also carried an undisclosed *Nestin-TVA* transgene (64). This transgene expresses a quail derived cDNA encoding the cell surface receptor for subgroup A avian leukosis virus in neural and glial stem and progenitor cells. The presence of the *Nestin-TVA* transgene did not materially affect the results or interpretations of experiments or the response of NRAS^Q61R^-driven melanoma to RMC-7977.

In the *TBC* and *TNC* genetically engineered mice, melanoma was initiated by 4-hydroxytamoxifen (4-HT) mediated activation of CreER^T2^ in melanocytes by topical administration of 1.5μl of 20μM (z)-4-hydroxytamoxifen (4-HT; H7904, Sigma-Aldrich) on two consecutive days in mice of both sexes, aged between 1-2 days (neonatal) – 6 months (adult). One day following 4-HT treatment (day 3), mice were irradiated with ultraviolet (UV)-B light at 3000J/m^2^. Mice were then monitored for melanomagenesis by visual inspection at least twice weekly. Melanoma volume was calculated as V=(length × width^2^)/2, where length is the greatest longitudinal diameter and width is the greatest transverse diameter, as measured by digital calipers.

For xenograft studies, 10^6^ OSUMMER or Mel9 cells **(Suppl. Table S2)** were resuspended in 100µl of sterile matrigel and subcutaneously engrafted into both flanks of Glowing Head (24) or NOD/SCID (Charles River) mice respectively. MTG2 or PDX-4872 **(Suppl. Table S2)** melanoma patient-derived xenografts (MPDX) were engrafted into a single flank of NOD/SCID mice (Charles River). The human MPDX models in Figs. 2G & H were generated using fresh tumor fragments obtained from Huntsman Cancer Hospital with written informed consent from patients in accordance with protocols approved by the Hospital Institutional Ethical Committee (IEC). Tumor fragments were subcutaneously serially passaged in immunodeficient mice and cryopreserved for future use. Recovered tumor fragments were implanted into the right flanks of immunodeficient mice; treatment started when the average tumor volume reached 100–300 mm^3^. Tumor outgrowth was monitored by digital caliper measurements over time. Tumors were allowed to grow in mice either for a pre-specified amount of time or until they reached endpoint based on the Ullmann-Culleré body conditioning score as specified in our approved IACUC protocol (31).

### *In vivo* preclinical therapeutic studies

RMC-7977 was synthesized according to a previously described protocol (9). For *in vivo* studies, RMC-7977 was prepared in a vehicle of 10%(v/v) DMSO (#D8418, Sigma-Aldrich), 20%(v/v), Polyethyleneglycol-400 (PEG400, MedChemExpress), 10%(v/v) Solutol HS 15 (Kolliphor HS 15, #42966, Sigma Aldrich), 60%(v/v) water, and administered at a dose of 10 mg/kg in 70-150μl volume by oral gavage. The same vehicle formulation was used for control groups. Tumors were harvested 2 hours after the final dose was administered. For cutaneous melanoma and xenografted tumor treatment studies, mice were enrolled onto vehicle or RMC-7977 treatment in a pseudo-randomized manner when exhibiting a cutaneous melanoma of ∼250 mm^3^. Human PDX studies in Figs. 2G & H were conducted at the following CROs: Pharmaron (Beijing, China), Champions Oncology (Hackensack, NJ), and Charles River Laboratories (Freiburg, Germany). All cell line derived xenograft (CDX)/patient derived xenograft (PDX) mouse studies and procedures related to animal handling, care and treatment complied with all applicable regulations and guidelines of Institutional Animal Care and Use Committee (IACUC) at each facility with their approvals. Female BALB/c nude, NOD SCID, NMRI nu/nu, and athymic Nude mice 6-12 weeks old were used. Animal vendors include Beijing AniKeeper Biotech Co. Ltd., Envigo, and Charles River Laboratories.

### Targeted Sequencing

Genomic DNA (gDNA) was extracted from cell lines using the DNeasy Blood&Tissue kit (Qiagen) and eluted in water. Plasmids were extracted from bacterial cell pellets using the QIAprep Spin Miniprep Kit (Qiagen). Complementary DNA (cDNA) was prepared from RNA using the High-Capacity cDNA Reverse Transcription Kit (Thermo). DNA was amplified by PCR at specific loci using primers listed in **Suppl. Table S10.** PCR products were analyzed using 3%(w/v) agarose gels and purified using the Zymoclean Gel DNA Recovery Kit (Zymo Research). Sanger sequencing was performed on gel extracted PCR products or purified plasmids by the HSC DNA Sequencing Core at the University of Utah.

### Whole exome sequencing

Genomic DNA (gDNA) was extracted from pellets of cultured melanoma cells using the Qiagen DNeasy Blood & Tissue kit. Exome libraries were prepared using the: 1. IDT xGEN Human Exome v2 with Nextera Flex Library Prep (4-plex enrichment) for human cell line samples or; 2. Twist Bioscience Mouse Exome panel. Subsequent Illumina sequencing employed the NovaSeq X Reagent Kit _150×150 bp and was performed by the HCI High-Throughput Genomics shared with DNA sequence analysis assisted by the HCI Bioinformatics Analysis shared resource. 400M read-pairs were split up between 6 mouse cell line samples (OSUMMER) or 5 human cell line samples (PDX4872). Mutations were called using the treatment-naive matched parental cell line as reference (PDX-4872). Exome sequences of parental OSUMMER.1 and OSUMMER.10 cells were previously published: accession numbers SRX20082359 and SRX20082371) (23).

### CRISPR/CAS9-mediated genome editing

Genome editing was performed in OSUMMER.1 cells. Alt-R™ S.p. CAS9 Nuclease V3, 100µg (#1081058), tracrRNA, and guides were purchased from Integrated DNA Technologies (IDT) and prepared according to the manufacturer’s instructions. To validate the CRISPR knockout, gene-edited sequences were PCR amplified and subjected to DNA sequencing. Primers and targeting sgRNAs were purchased from Integrated DNA Technologies (IDT) (**Suppl. Table S11).**

### RNA extraction, cDNA preparation, and quantitative RT-PCR

RNA was extracted from cell lines using the RNeasy Mini Kit (Qiagen 74106) following the manufacturer’s protocol. cDNA was prepared from 0.5-1μg of RNA using the Applied Biosystems High-Capacity cDNA Reverse Transcription Kit (ThermoFisher 4368814) according to the manufacturer’s protocol. cDNA was then used for qRT-PCR analysis using TaqMan Gene-Expression Master Mix (ThermoFisher 4369016). TaqMan primer probes specific to mouse *Ppia* (Mm03024003_g1 (FAM-MGB), Mm02342429_g1 (FAM-MGB), Mm02342430_g1 (FAM-MGB), Mm03302254_g1 (FAM-MGB) ThermoFisher) and mouse *Gapdh* (Mm99999915_g1, VIC-MGB_PL ThermoFisher) were duplexed to detect the levels from each sample in triplicates using a 10μl final reaction volume in a 384-well clear optical reaction plate (Applied Biosystems 4309849). Thermocycle analysis was performed using QuantStudio 6 Flex Real Time PCR System using the preprogrammed thermocycle run settings for comparative qPCR (Hold stage – step 1 50 °C for 2 minutes, step 2 95 °C for 10 minutes PCR stage – step 1 95 °C for 15 seconds, step 2 60 °C for 1 minute) for qPCR, Ct values of *Ppia* and *Gapdh*. *Ppia* Ct value was normalized to *Gapdh*, and then the mean relative mRNA expression levels of each group were normalized to the wild-type control group. Values were plotted as relative change in mRNA expression compared with wildtype. Means ± SEM were shown.

### Quantitative PCR (qPCR) on genomic DNA (gDNA)

Genomic DNA (gDNA) was extracted from cell lines using the DNeasy Blood&Tissue Kit (Qiagen 69504). The zygosity of *Ppia* exons was determined using quantitative PCR (qPCR) analysis with 25µg of gDNA, 20µM primer, and 10µM probe. The probe contained a 5’FAM fluorophore with a ZEN and 3’ Iowa Black FQ (IBFQ) quencher. gDNA was amplified by PCR at specific loci using primers listed in **Suppl. Table S12.**

### Site directed mutagenesis

A plasmid encoding human Cyclophilin A (CYPA) expressed from the SFFV promoter and puromycin resistance from the CYPA promoter (pScalps-*huPPIA_WT*-Puro) was kindly provided by Dr. Jeremy Luban (U. Massachusetts Medical School). Point mutations in the CYPA cDNA: 1. T407G encoding CYPA^M136R^; 2. T186G encoding CYPA^C62W^; 3. C109T encoding CYPA^R37C^ and; 4. C376-377AT encoding the CYPA^H126HX^ frameshift alteration were engineered in pScalps-*huPPIA_WT*-Puro. A plasmid expressing human wild-type MEK1 (MAP2K1) and puromycin resistance (pBABEpuro-HA-MEK1) was purchased from Addgene (plasmid #53195). The *Map2k1^C371T^* missense mutation was engineered in pBABEpuro-HA-MEK1 to encode MEK1^P124L^. Mutations were introduced by site-directed mutagenesis using non-overlapping primers using the New England Biolabs (NEB) Q5 SDM protocol and were designed by the NEBaseChanger program (primer sequences see **Suppl. Table S13**). DNA was transformed into 5-α chemically competent bacteria (NEB), and bacterial DNA was extracted from single colonies using the Qiagen Miniprep kit. All site-directed alterations in the *PPIA or MAP2K1* cDNA were confirmed by DNA sequencing.

### Immunohistochemical staining

Haematoxylin and eosin (H&E) staining and immunohistochemistry (IHC) were performed on 4µm formalin-fixed paraffin-embedded (FFPE) sections that had previously been baked at 60°C for two hours. FFPE sections were deparaffinized using xylene and epitope retrieval using Dako target retrieval solution pH6 (#S236984-2, Agilent) for 30 min at 95°C. Sections were rinsed with Milli-Q ultrapure water before endogenous peroxidase blocking using a BLOXALL endogenous blocking solution (#SP-6000-100, Vector Laboratories) for 10 min. Sections were rinsed with Tris-Buffered Saline with 0.1%(v/v) Tween 20 (TBST) before applying a 2.5%(v/v) normal horse serum blocking solution (#S-2012-50, Vector Labs) for 30 minutes at room temperature. Sections were then rinsed with TBST before application of primary antibodies at an optimized dilution (ERK1/2 phospho-T202/Y204 (CST#4370, 1:600); Cyclophilin A (ProteinTech #10720-1-AP, 1:150); Ki67 (D3B5; CST#12202, 1:400); S1004A (D9F9D; CST#13018, 1:500), for 1 hour at room temperature. Sections were then rinsed with TBST and incubated for 30 minutes with an anti-rabbit ImmPRESS Polymer Reagents (VectorLaboratories). Sections were rinsed with wash buffer and visualized using ImmPACT 3,3′-diaminobenzidine (DAB) substrate kit (SK-4105, Vector Laboratories). Finally, sections were washed in tap water and counterstained with hematoxylin. (#HHS32-1L, Millipore Sigma).

### Immunoblot Analysis

Cells were trypsinized, washed in PBS, and resuspended in RIPA buffer supplemented with 1x Protease and Phosphatase Inhibitor Cocktail (Thermo Scientific). Tumor chunks were dissociated and made into protein lysates using RIPA buffer supplemented with protease inhibitors (Thermo) as per manufacturer guidelines. Lysates were electrophoresed on duplicate 4-12%(w/v) gradient SDS-PAGE gels and transferred onto PVDF membranes at 20V for 7 minutes using the Invitrogen iBlot2 system. PVDF membranes were probed overnight at 4°C with primary antibodies as follows: pT202/Y204-ERK1/2 (D13.14.4E, CST#4370 1:500); pS473-AKT (D9E, CST#4060 1:500); pan-AKT (40D4, CST#2920 1:500); pS235/236-Ribosomal Protein S6 (D57.2.2E, CST#4858 1:500); pS65-4EBP1 (CST#9451 1:500); pT246-PRAS40 (CST#2997 1:500); Cyclophilin A C-terminus (CST#2175 1:500, reactivity: mouse or human); Cyclophilin A N-terminus (CST#51418 1:500, reactivity: mouse); PTEN (D4.3) XP (CST#9188 1:500), HA-tag (CST#3724 1:1000), GAPDH (CST#97166 1:1000), β-Actin (CST#3700 1:1000). One of the two duplicate blots was generally left uncut and incubated with a primary antibody cocktail that is known to generate distinct bands at known molecular weights (e.g. pAKT, tAKT, pS6, p4EBP1, ACTIN). The other duplicate blot was generally cut into smaller pieces and incubated with the remaining antibodies of interest. Primary antibody binding was detected using secondary antibodies including fluorescent goat anti-Rabbit IRDye 800 or goat anti-Mouse IRDye 680 (LI-COR Biosciences), which were applied simultaneously as a secondary antibody cocktail, allowing for visualization of proteins at the same molecular weight in the red or green channels respectively. The Invitrogen SeeBlue Plus2 Prestained Protein Standard was used as molecular weight markers. Immunofluorescent signals were visualized using a LI-COR Odyssey CLx imaging system and Image Studio V5.2 software (LI-COR Biosciences).

### Statistical analysis and data visualization

Statistical analysis and graph plotting was conducted using GraphPad Prism 10 (La Jolla, USA). All comparisons made, and statistical tests used are described in each figure legend. IncuCyte S3 Software 2022 was used to calculate standard deviations and standard errors from cell confluency. Prism 10 GraphPad (LaJolla, USA) was used to generate graphs from processed IncuCyte data. Cell Confluency Assays were performed in triplicate wells (biological replicates) per cell line and per treatment group for each experiment.

### Data Availability

Data used to generate analyses and visualization in this publication are available within the article and its Supplementary data files.

## ACKNOWLEDGEMENTS

We thank the members of the McMahon and Kinsey laboratories for advice, guidance and support on this project. We are grateful to, and acknowledge: 1. Dr. David Lum and colleagues of the HCI Preclinical Research Shared Resource for assistance with implantation of PDXs and drug dosing of experimental mice; 2. Dr. Allie Grossmann for histopathological analysis of mouse melanoma specimens; 3. Drs. Kenneth Boucher and Ann Chen for advice and guidance on statistical analyses; 4. The HCI High-Throughput Genomics and Bioinformatics Shared Resources for assistance with Next Generation Sequencing and analysis, respectively; 5. The University of Utah Health Sciences Center (UU-HSC) DNA Sequencing core for assistance with Sanger sequencing, cell line authentication, and digital PCR experiments; 6. The HCI Immuno-Oncology Initiative (IOI) initiative for assistance with tumor dissociation and cell line preparation from biospecimens; 7. The ARUP Research Histology Core for preparation of H&E and unstained slides; 8. The UU-HSC Flow Cytometry Core Facility for assistance with single cell sorting; 9. The UU Office of Comparative Medicine for mouse veterinary care and husbandry; 10. Dr. Christin Burd (Ohio State University) for access to the OSUMMER *NRAS*-driven melanoma cell lines and the *Nras^LSL-Q61R^*mouse strain for the generation of the *TNC* GEM model of NRAS-driven melanoma; 11. Dr. Doug Grossman for advice, guidance, and access to a UV exposure chamber; 12. Dr. Jeremy Luban (University of Massachusetts Medical Center) for the gift of the pScalps-CYPA construct; and 13. Dr. Keiran Smalley (Moffitt Cancer Center) for ongoing collegial discussions. This research was funded by: 1. A sponsored research agreement from Revolution Medicines; 2. The HCI Melanoma Disease Center; 3. The HCI Comprehensive Cancer Support grant (P30 CA042014); 4. An HCI’s GEMS T32 to K.O.T; 5. The University of Utah, Department of Dermatology supporting D.C.D., 6. The Huntsman Cancer Foundation; 7. The Beesley Family Foundation; and 8. An NCI R01 (CA176829) to MM.

## SUPPLEMENTARY FIGURE LEGENDS

**SUPPLEMENTARY FIGURE S1:**
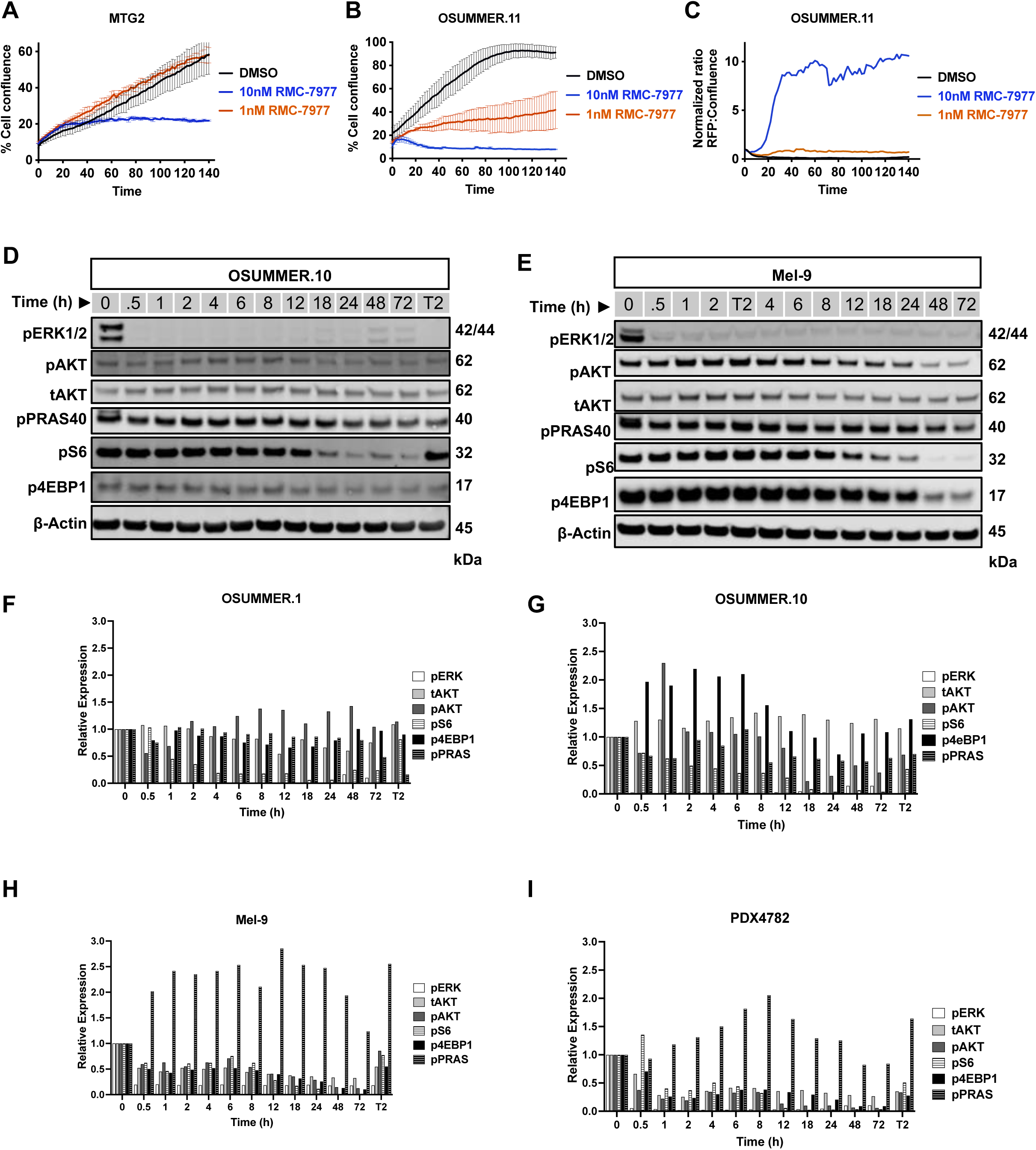
(**A-C**) Cell confluency or cytotoxicity assays under pharmacological inhibition. **(A)** Human NRAS-driven melanoma cell line MTG2 (NRAS^Q61R^) or **(B, C)** mouse NRAS-driven melanoma cell line OSUMMER.11 (NRAS^Q61R^) were treated with vehicle (DMSO) or RMC-7977 at indicated concentrations and analyzed for cell confluence **(A-B)** or cytotoxicity by CytoxRed reagent uptake **(C)** using an Incucyte Live-Cell Analysis System over time. All Incucyte experiments were performed on three separate occasions. Error bars indicate SD of triplicate wells. **(D-E)** Immunoblot analysis of cell lysates prepared from mouse melanoma cell lines OSUMMER.10 (NRAS^G13R^; **D)** or Mel9 (NRAS^Q61R^; **E)** treated with DMSO or 80nM RMC-7977 over a time course of 0.5-72 hours. A two-hour 100nM trametinib control treatment (T2) was included in each run for complete suppression of pERK1/2. Lysates were analyzed for the phosphorylation (p) or total (t) abundance of ERK1/2, AKT, PRAS40, S6 ribosomal protein, 4EBP1, or β-actin, as indicated. **(F-I)** Protein bands quantification of OSUMMER.1, OSUMMER.10, PDX4872, and Mel9 immunoblots depicted in **Main Fig.1 I-J,** and **Suppl. Fig. S1 D-E** over the indicated time course using Image Studio V5.2 software (LI-COR Biosciences).

**SUPPLEMENTARY FIGURE S2:**
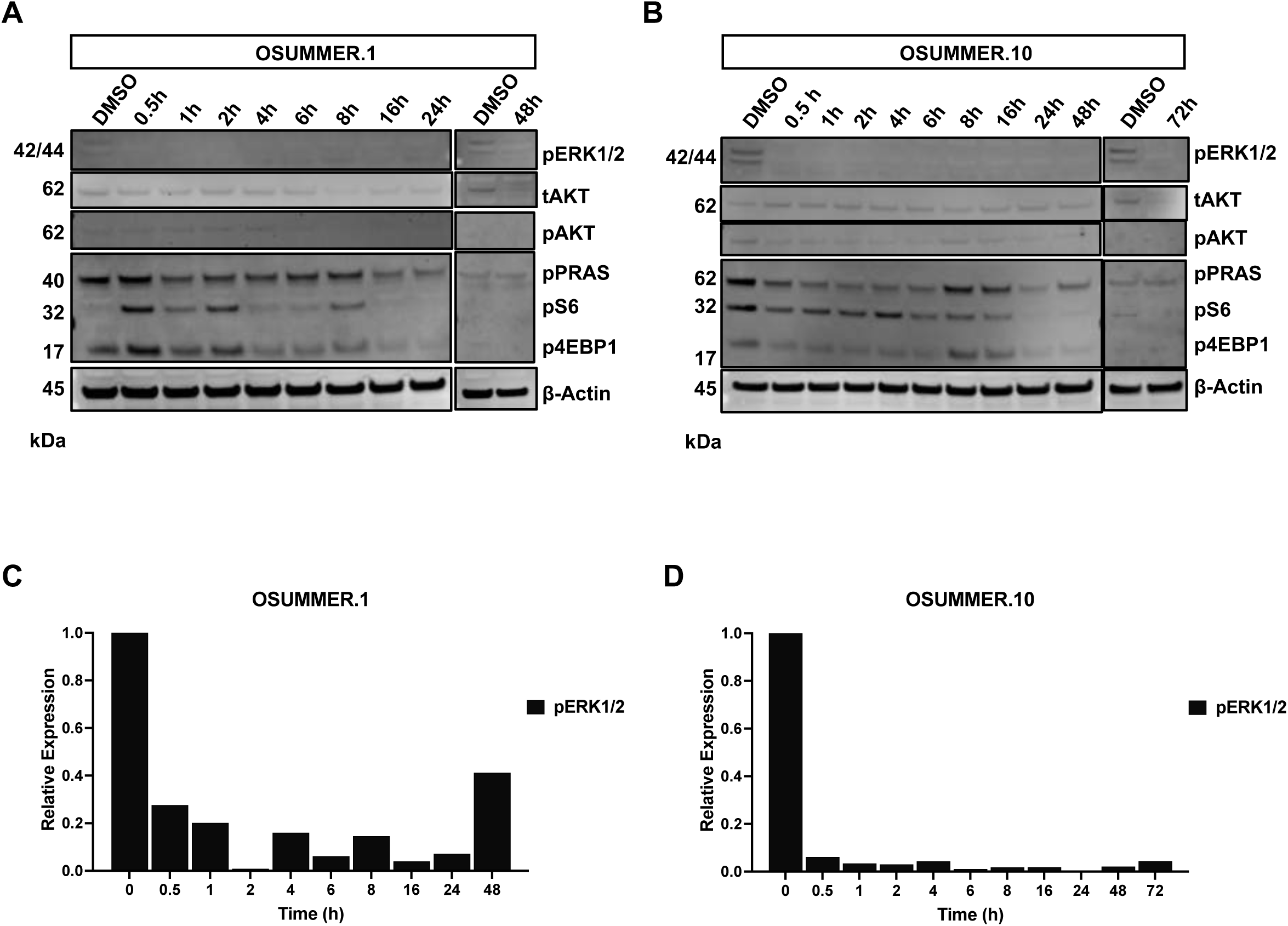
(**A-B**) Immunoblot analysis of cell lysates prepared from mouse melanoma cell lines OSUMMER.1 (NRAS^Q61R^; **A)** or OSUMMER.10 (NRAS^G13R^; **B)** treated with DMSO or 25nM RMC-7977 over a time course of 0.5-48 (O.1) or 0.5-72 (O.10) hours. Lysates were analyzed for the phosphorylation (p) or total (t) abundance of ERK1/2, AKT, PRAS40, S6 ribosomal protein, 4EBP1, or β-actin, as indicated. **(C-D)** Protein bands quantification of OSUMMER.1 **(C)** or OSUMMER.10 **(D)** immunoblots depicted in **(A, B)** over the indicated time course using Image Studio V5.2 software (LI-COR Biosciences).

**SUPPLEMENTARY FIGURE S3:**
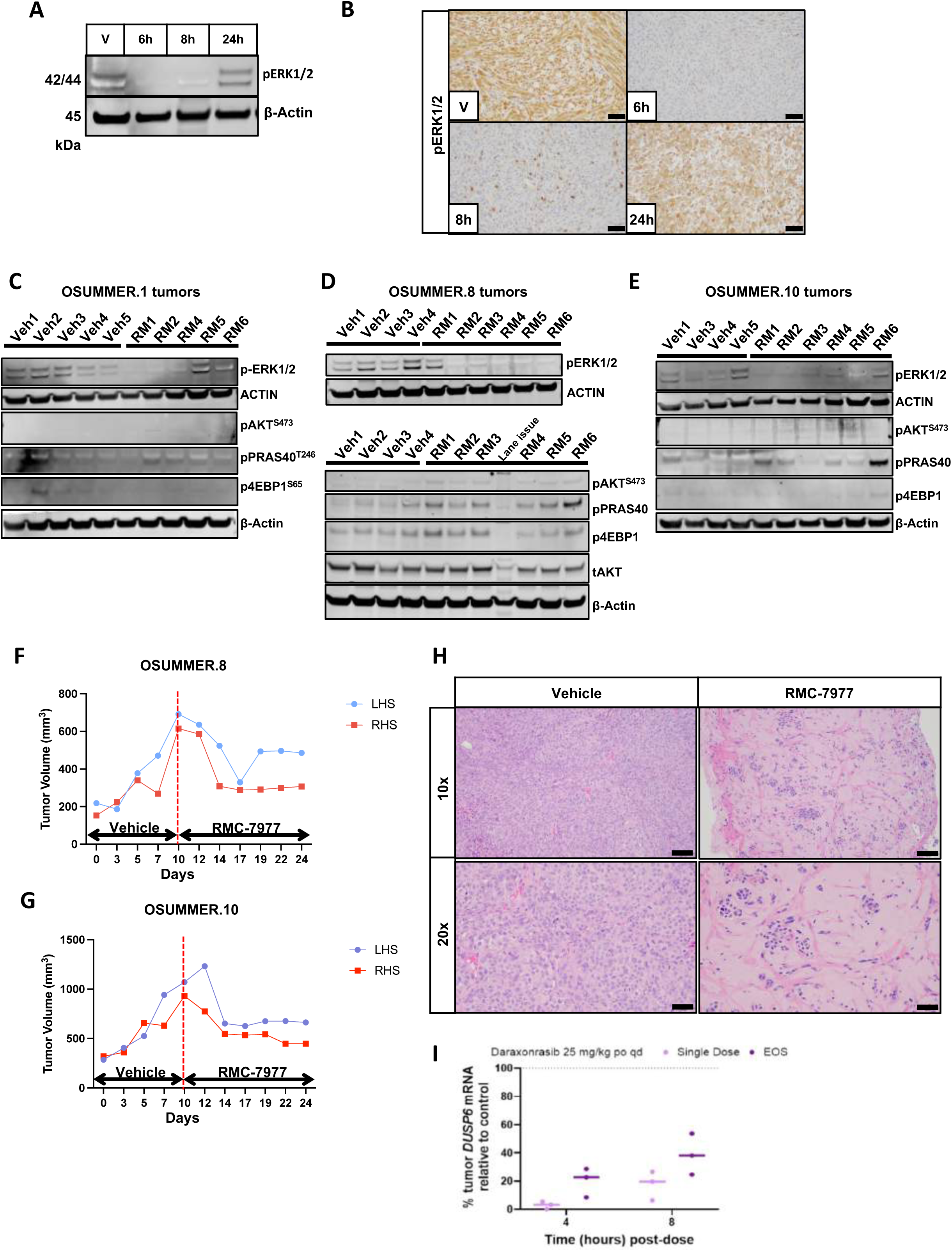
(**A-B**) Pharmacodynamics of RMC-7977 in preclinical OSUMMER.1 allograft model. Mice were injected with 10^6^ OSUMMER.1 (NRAS^Q61R^) cells on each flank and tumors were matured to ∼300 mm^3^. Mice were treated with a single dose of 10 mg/kg of RMC-7977, tumors were harvested, cut in half, and one half was made into protein lysates at 6h, 8h, or 24h after dosing, along with a vehicle treated mouse tumor (V). Lysates were analyzed for the pERK1/2, or β-actin, as indicated **(A)**. The other half of the tumors was formalin-fixed, paraffin-embedded, and IHC for pERK1/2 was performed on representative sections **(B).** Scale bar in lower right corner represents 50uM in all images. **(C-E)** Immunoblot analysis of cell lysates prepared from mouse melanoma allograft tumors OSUMMER.1 (NRAS^Q61R^; **C)**, OSUMMER.8 (NRAS^Q61K^; **D)**, or OSUMMER.10 (NRAS^G13R^; **E)** of mice that were treated with vehicle or 10 mg/kg RMC-7977 until experimental endpoint at 28-32 days depicted in Main Fig. 2A-C). Lysates were analyzed for the phosphorylation (p) or total (t) abundance of ERK1/2, AKT, PRAS40, S6 ribosomal protein, 4EBP1, or β-actin, as indicated. **(F-G)** Mouse tumor volume graphs of two mice allografted with either OSUMMER.8 (NRAS^Q61K^, **F**) or OSUMMER.10 (NRAS^G13R^, **G**) in each flank (left-hand side; LHS or right-hand side; RHS). Mice were treated with vehicle for 10 days until a tumor volume of 600-1000 mm^3^ had been reached (red dotted line), and with 10 mg/kg RMC-7977 thereafter until experimental endpoint at 24 days. **(H)** Representative H&E images of sections of Mel9 tumors grown in NOD/SCID mice showing hyalinizing stroma as a treatment effect of RMC-7977 at endpoint (18 days of treatment) at 10x and 20x as indicated. Scale bar in lower right corner represents 100uM in 10x and 50uM in 20x images. **(I)** Percent tumor DUSP6 messenger RNA (mRNA) relative to control at 4 or 8 hours post single dose of 25 mg/kg daraxonrasib (pink) or at end-of-study during apparent on-treatment relapse (EOS; purple).

**SUPPLEMENTARY FIGURE S4:**
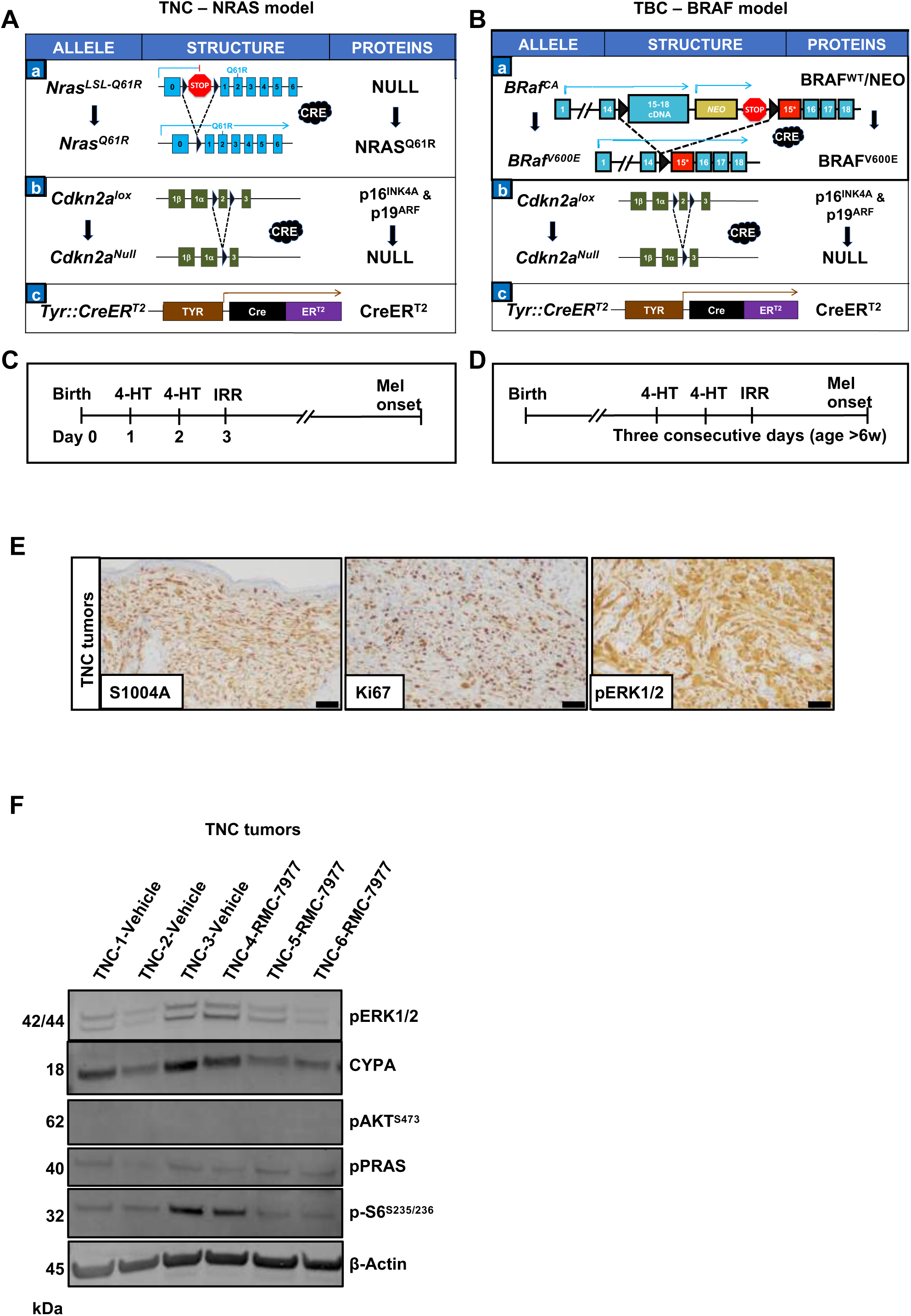
(**A-B**) Schematic of *TyrCreER^T2^ Nras^LSL-Q61R/Q61R^ Cdkn2a^flox/flox^ (TNC)* or *Tyr::CreER^T2^ Braf^CA/+^ Cdkn2a^flox/flox^ (TBC)* mouse alleles detailing the genetic alleles of **(A, a)** *NRAS^LSL-Q61R^* allowing for Cre-mediated expression of NRAS^Q61R^, **(A, b)** *Cdkn2a^lox^*allowing for Cre-mediated silencing of the p16^INK4A^ and p19^ARF^ tumor suppressors, and **(A, c)** *Tyr::CreER^T2^* providing for conditional, melanocyte-specific activation of Cre recombinase; **(B, a)** *Braf^CA^*allowing for Cre-mediated expression of BRAF^CA^ in any target tissue of the mouse, **(B, b)** *Cdkn2a^lox^* and **(B, c)** *Tyr::CreER^T2^* as described above. **(C-D)** Schematic of the 4-hydroxytamoxifen (4-HT) and UVB irradiation (IRR) regimen for *TNC* and *TBC* models to initiate Cre recombination and tumor growth in neonates (**C**) or in adults (**D**; > 6 weeks of age; detailed in Methods). **(E)** IHC for S1004A, Ki67, and pERK1/2 on representative sections from untreated *TyrCreER^T2^; Nras^LSL-Q61R/Q61R^; Cdkn2a^flox/flox^ (TNC)* tumors of mice that were induced as adults by administration of 4-HT and subsequent UV-irradiation. Scale bar in lower right corner represents 50uM in all three images. **(F)** Immunoblot of lysates from *TNC* mouse tumors that were induced by administration of 4-hydroxytamoxifen-(4-HT) and subsequent UV-irradiation. Mice were treated with vehicle (n=3) or 10 mg/kg RMC-7977 (n=3) for 28 days until resistance had become evident by increase in tumor volume. Mice received their last dose of vehicle or RMC-7977 two hours prior to tumor harvest. Lysates were analyzed for pERK1/2, CYPA, pAKT, pPRAS40, pS6 ribosomal protein, p4EBP1, or β-actin, as indicated.

**SUPPLEMENTARY FIGURE S5:**
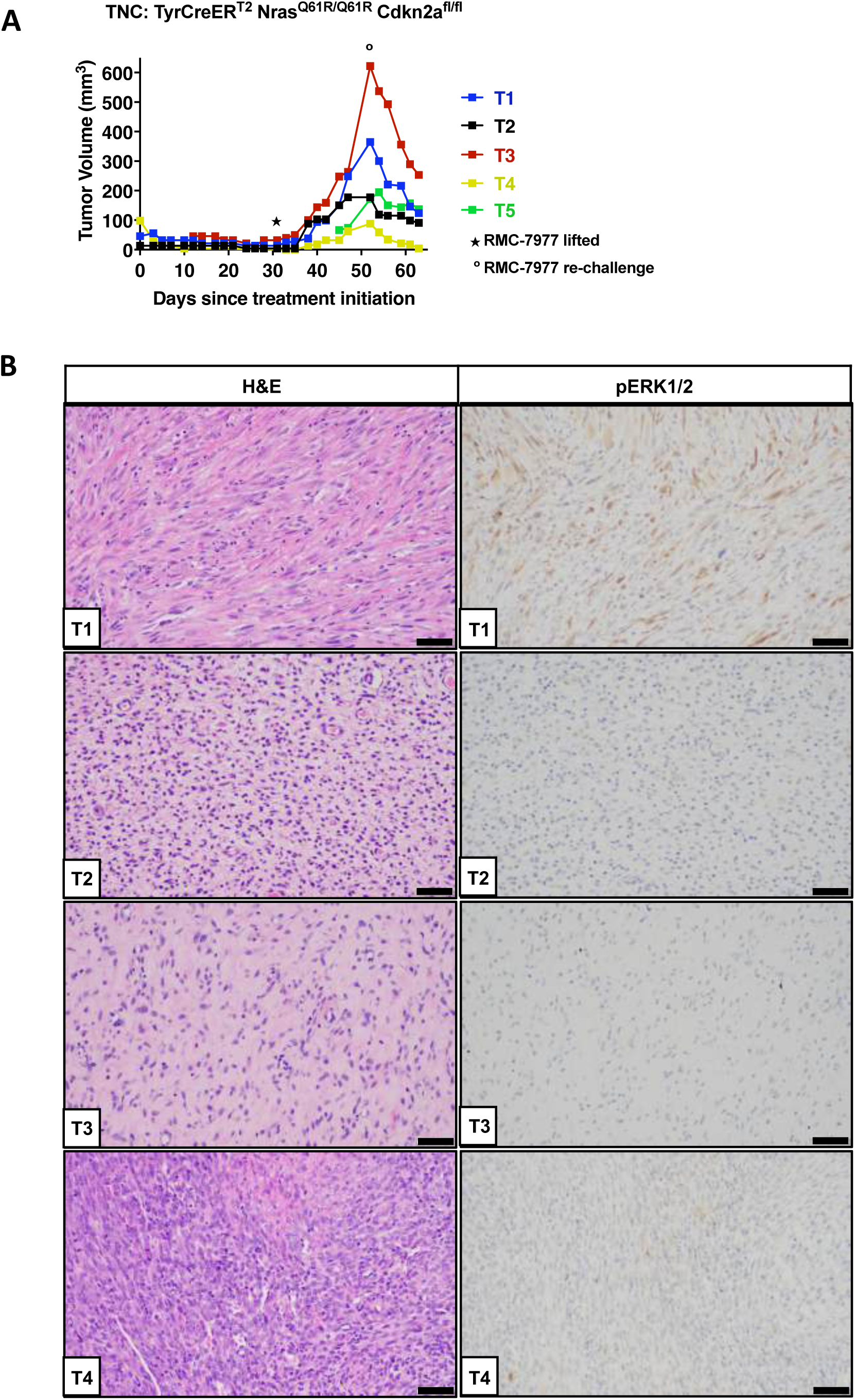
(**A**) Tumor volume graph of five tumors from a single *TyrCreER^T2^ Nras^LSL-Q61R/Q61R^ Cdkn2a^flox/flox^* (*TNC*) mouse induced as a neonate, and treated with 10 mg/kg RMC-7977 for 30 days. Star (11) at 30 days indicates time point where RMC-7977 was discontinued. Circle (ο) at 50 days indicates re-challenge with 10 mg/kg RMC-7977 daily dosing until end of experiment. **(B)** H&E and IHC for pERK1/2 on four out of the five *TNC* tumors. Mice received a final dose of RMC-7977 two hours prior to tumor harvest. Scale bar in lower right corner represents 50μM in all images.

**SUPPLEMENTARY FIGURE S6:**
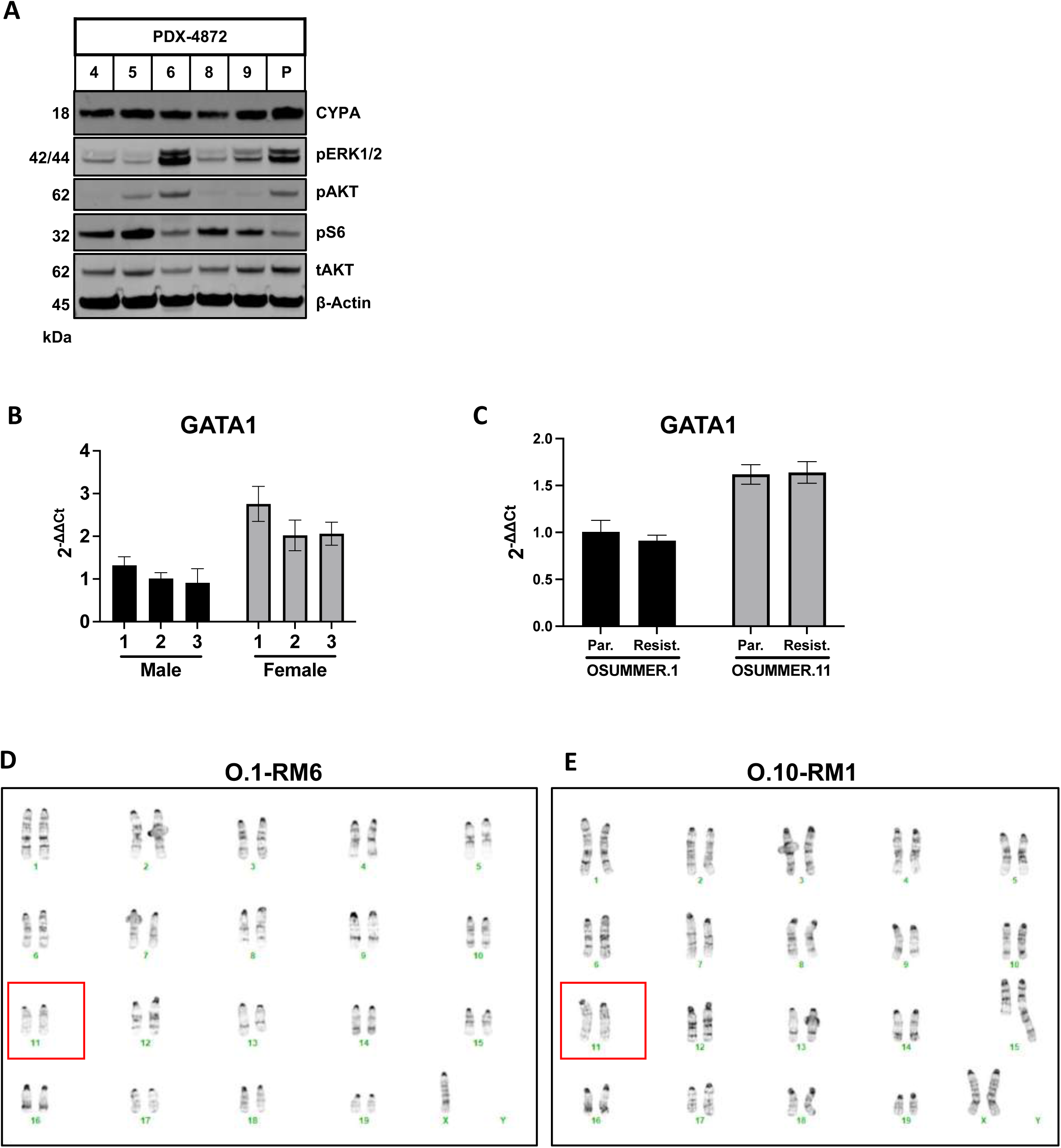
(**A**) Immunoblot analysis of cell lysates prepared from human PDX-derived cell line parental PDX-4872 (P) grown in the absence of RMC-7977, or RMC_7977^R^ clones 4, 5, 6, 8, 9 grown in the presence of 100nM RMC-7977. Lysates were analyzed for CYPA, pERK1/2, pAKT, pS6 ribosomal protein, or β-actin, as indicated. **(B-C)** Quantitative PCR (qPCR) for X-chromosome-linked gene GATA1 on genomic DNA from three male (black bars) vs. three female mice (gray bars; **B**) or for X-chromosome-linked gene GATA1 on genomic DNA from male OSUMMER.1 parental (Par.) or resistant (Resist.) cells (black bars) vs. female OSUMMER.10 parental or O.10^RM1^ RMC-7977 resistant cells (gray bars; **C**). Y-axis (2^ΔΔCt) represents the difference in Ct values between the target gene and the reference gene (fold change in gene expression), calculated across the different experimental groups. **(D-E)** Karyotyping of O.1^RM6^ or O.10^RM1^ indicating an intact chromosome 11 (red box) in both cell lines.

**SUPPLEMENTARY FIGURE S7:**
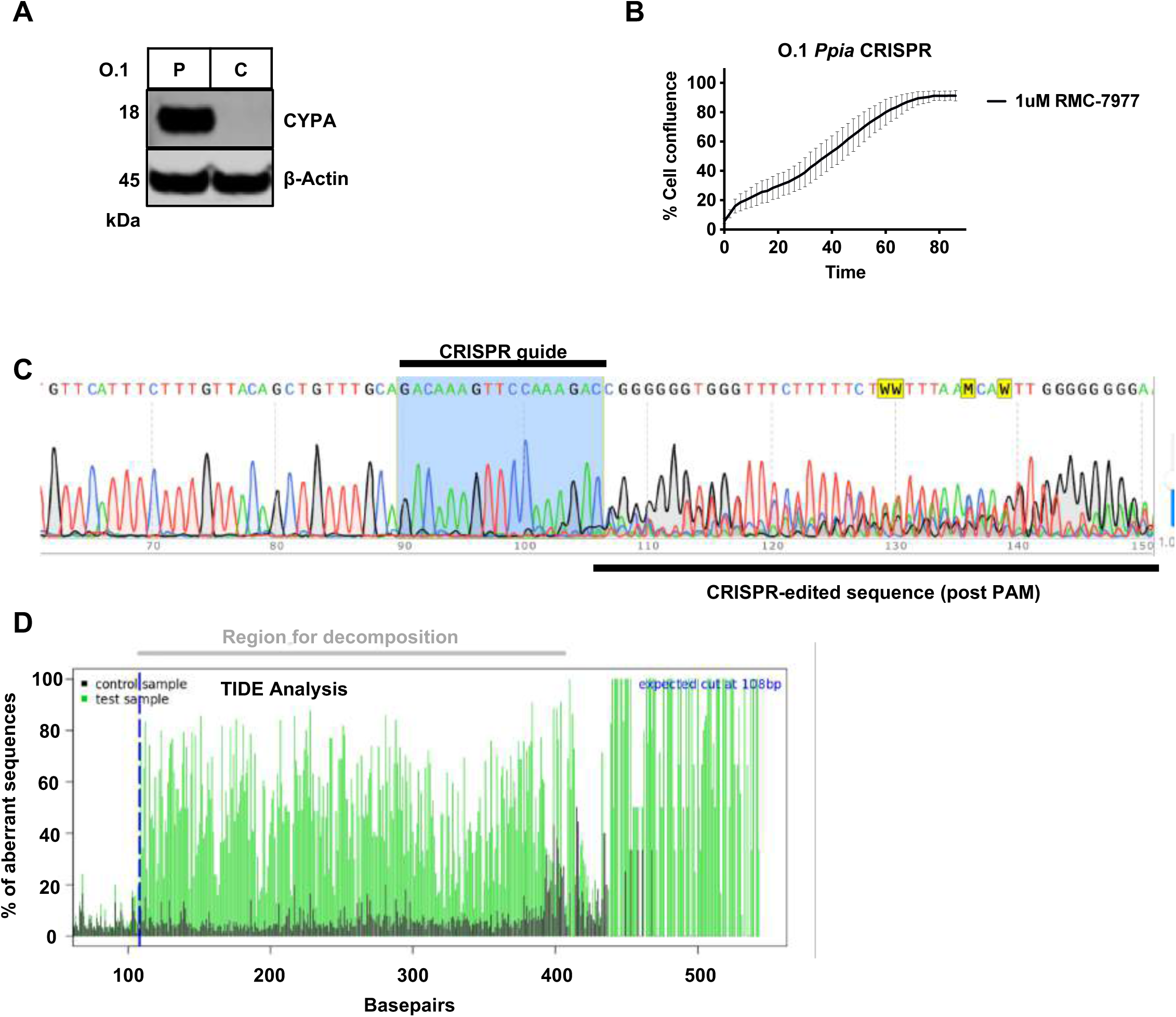
(**A**) Immunoblot for total CYPA protein, pERK1/2, or β-actin on lysates from parental OSUMMER.1 (P) versus O.1 *Ppia* CRISPR knockout cells (C). **(B)** Cell confluency assay of O.1-*Ppia*-CRISPR under 1μM RMC-7977 as analyzed by Incucyte Live-Cell Analysis. Graph is representative of n=3 independent experiments. Error bars represent standard deviation (SD) of triplicate technical replicates. **(C)** Sanger sequencing of pooled population of O.1-*Ppia*-CRISPR showing sgRNA (CRISPR guideRNA) alignment and CRISPR-edited sequence downstream of the PAM sequence. **(D)** TIDE analysis showing the region of decomposition after the CRISPR guide RNA binding site in Control (unedited sequence of OSUMMER.1) versus Test sample (O.1 *Ppia* CRISPR pool).

**SUPPLEMENTARY FIGURE S8:**
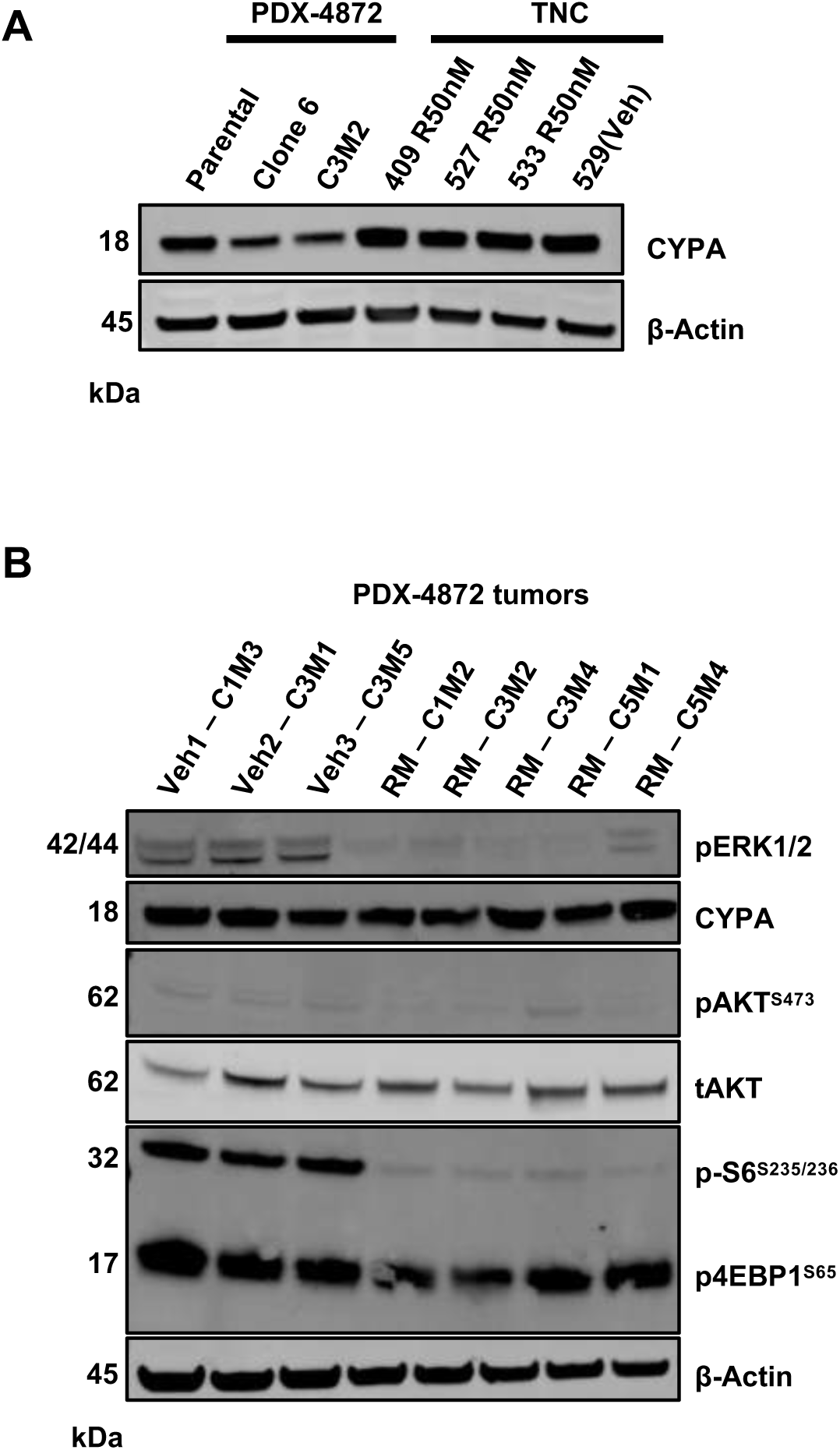
(**A**) Immunoblot analysis of cell lysates prepared from human PDX-derived cell lines PDX-4872-Parental, or RMC-7977^R^ cell lines PDX-4872-clone 6 or –C3M2 (lanes 1-3), or from RMC-7977^R^ *TNC (TyrCreER^T2^; Nras^LSL-Q61R/Q61R^; Cdkn2a^flox/flox^*) cell lines #409, #527, #533, or #529-Veh (lanes 4-7). Lysates were analyzed for CYPA expression or β-actin, as indicated. **(B)** Immunoblot analysis of cell lysates prepared from PDX tumors of PDX-4872 of mice that were treated with vehicle or 10 mg/kg RMC-7977 until experimental endpoint at 28 days depicted in Main Fig. 2G. Lysates were analyzed for the phosphorylation (p) or total (t) abundance of ERK1/2, CYPA, AKT, PRAS40, S6 ribosomal protein, 4EBP1, or β-actin, as indicated.

**SUPPLEMENTARY FIGURE S9:**
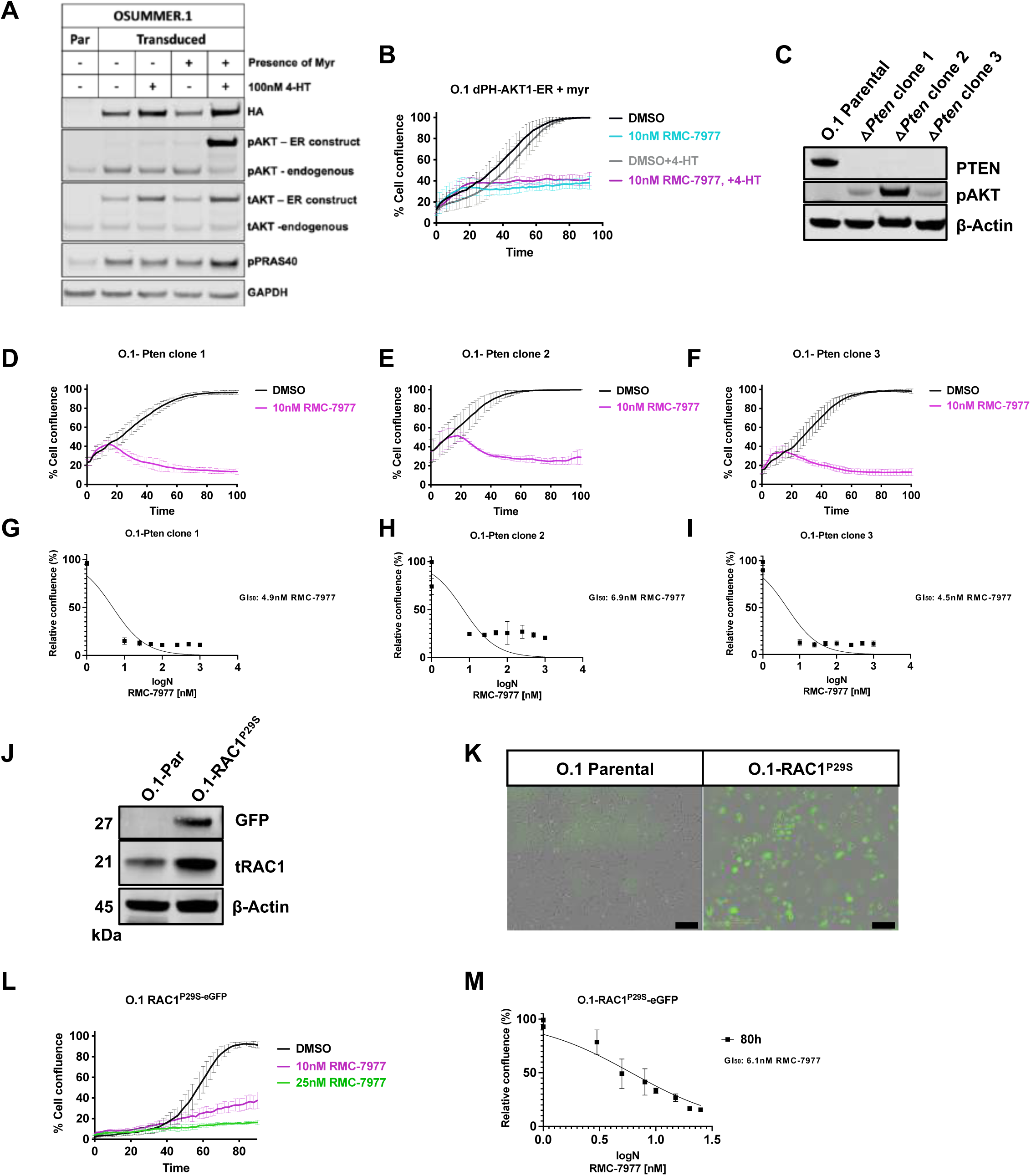
(**A**) Immunoblot of OSUMMER.1-Parental (Par) vs. OSUMMER.1-derived cell line engineered to stably express a conditionally activated, myristoylated (myr)AKT1:ER construct in the absence or presence of 4-hydroxytamoxifen (4-HT) as indicated, probed for HA-tag of (Myr+/-) AKT1 construct, pAKT, tAKT, pPRAS40, or GAPDH (37,38). pAKT antibody detecting the estrogen receptor (ER)-tagged construct or endogenous pAKT, tAKT antibody detecting the ER-tagged construct or endogenous tAKT. **(B)** Incucyte proliferation assay measuring percent cell confluence over time in OSUMMER.1-Parental vs. OSUMMER.1-derived cell line stably expressing inducible myrAKT1:ER in the presence of 4-HT or 10nM RMC-7977 as indicated. Graph is representative of n=3 independent experiments. Error bars represent standard deviation (SD) of triplicate technical replicates. **(C)** Immunoblot for PTEN, pAKT or β-actin in OSUMMER.1-Parental vs. three separate OSUMMER.1-derived clones with CRISPR-mediated silencing of PTEN expression. **(D-F)** Incucyte proliferation assay measuring percent cell confluence over time in OSUMMER.1-*Pten^Δ^* knockout clones treated with DMSO or 10nM RMC-7977 as indicated. **(G-I)** Viability levels of OSUMMER.1-*Pten ^Δ^* clones treated with indicated logN concentrations of RMC-7977 in nM for 80 hours with GI_50_ values indicated in graphs. Graphs are representative of n=3 independent experiments. Error bars represent standard deviation (SD) of triplicate technical replicates. **(J)** Immunoblot for GFP, total RAC1, and β-actin in OSUMMER.1-Parental (Par) or O.1 cells that stably express an activating melanoma hotspot mutation, RAC1^P29S^. **(K)** GFP fluorescence of OSUMMER.1-Parental or O.1-RAC1^P29S^ cells. Scale bar in lower right corner represents 200μM in both images. **(L)** Incucyte proliferation assay measuring % cell confluence over time in OSUMMER.1-derived cell line stably expressing an activating melanoma hotspot mutation, RAC1^P29S^, treated with DMSO, 10nM or 25nM RMC-7977 as indicated. **(M)** Viability levels of O.1-RAC1^P29S^ treated with indicated logN concentrations of RMC-7977 in nM for 80 hours with GI_50_ value indicated in graph. Graphs are representative of n=3 independent experiments. Error bars represent standard deviation (SD) of triplicate technical replicates.

**SUPPLEMENTARY FIGURE S10:**
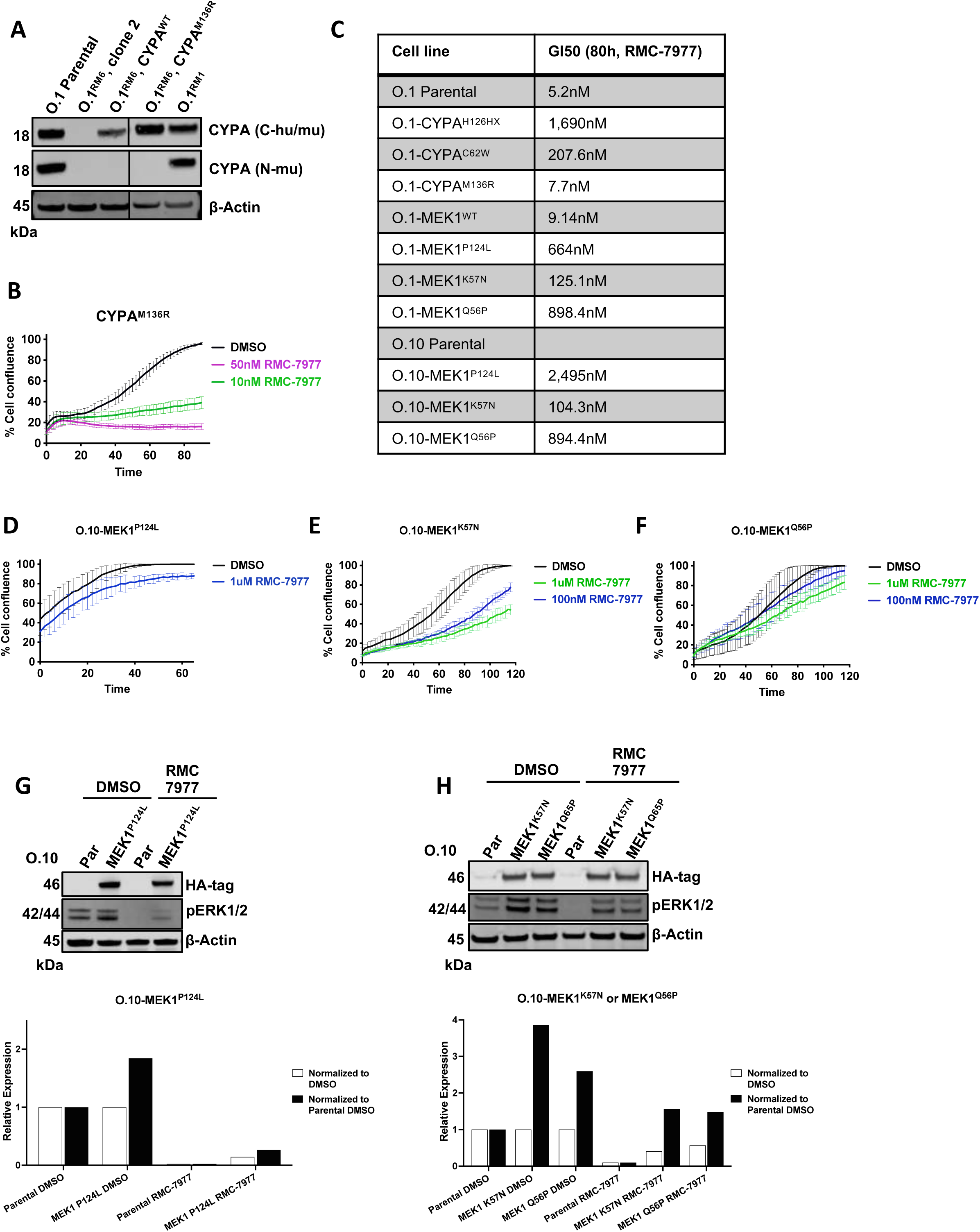
(**A**) Immunoblot for total CYPA protein using C– or N-terminus selective antibodies of the CYPA protein on lysates from parental OSUMMER.1, O.1^RM6^ clone 2, O.1^RM6^-CYPA^WT^, or O.1^RM6^-CYPA^M136R^. O.1^RM1^ is the original cell line that expressed a heterozygous copy of CYPA^M136R^ that was identified by WES. **(B)** Cell confluency assay of O.1^RM6^-CYPA^M136R^ treated with RMC-7977 as indicated and analyzed by Incucyte Live-Cell Analysis. Graph is representative of n=3 independent experiments. Error bars represent standard deviation (SD) of triplicate technical replicates. **(C)** 50% growth inhibition (GI_50_) values of cell lines tested for their viability levels under increasing concentration of RMC-7977 for 80 hours. GI_50_ values are representative of 2-3 independent experiments. **(D-F)** Cell confluency assays of OSUMMER.10 ectopically expressing MEK1^P124L^ **(D),** MEK1^K57N^ **(E),** or MEK1^Q56P^ **(F)** under concentrations of RMC-7977 as indicated and analyzed by Incucyte Live-Cell Analysis. Graphs are representative of n=3 independent experiments. Error bars represent standard deviation (SD) of triplicate technical replicates. **(G, H)** Immunoblots for HA-MEK1 fusion protein, pERK1/2 (quantification below panel), or β-actin on lysates from parental O.10 (Par), or O.10 ectopically expressing MEK1^P124L^ **(G),** or MEK1^K57N^ or MEK1^Q56P^ **(H)** in the absence or presence of 100nM RMC-7977 as indicated.

**Supplementary Table S1:** Cell lines used in Main Fig. 1A and 1B (Separate Excel File)

**Supplementary Table S2:** Cell lines.

| Cell line | Derived from | NRAS status | Source |
| --- | --- | --- | --- |
| OSUMMER.1 | Mouse; UV-induced TyCreER LSL-NRAS GEM model | Q61R | Millipore |
| OSUMMER.8 | Mouse; UV-induced TyCreER LSL-NRAS GEM model | Q61K | Millipore |
| OSUMMER.10 | Mouse; UV-induced TyCreER LSL-NRAS GEM model | G13R | Millipore |
| OSUMMER.11 | Mouse; UV-induced TyCreER LSL-NRAS GEM model | Q61R | Millipore |
| MTG2 | Human PDX-derived 2D melanoma cell line | Q61R | Huntsman Cancer Institute |
| Mel-9 | Human Biopsy-derived 2D melanoma cell line | Q61R | Huntsman Cancer Institute |
| PDX-4872 | Human PDX-derived 2D melanoma cell line | Q61K | Huntsman Cancer Institute |
| A375 | Human cutaneous melanoma | WT | ATCC |
| WM793 | Human cutaneous melanoma | WT | ATCC |
| HEK293T | Human embryonic kidney | WT | ATCC |

**Supplementary Table S3:** Patient-derived xenografts (PDXs)

| Patient-derived xenografts | NRAS status | Other known genetic alterations | Source |
| --- | --- | --- | --- |
| MTG2 (HCI-CM002) | Q61R | See PMID: 39627834 | Huntsman Cancer Institute |
| Mel-9 | Q61R | No additional sequencing information available | Huntsman Cancer Institute |
| PDX-4872 | Q61K | CRKL amplification, LYN amplification, MYC amplification, TERT promoter -146C>T; MS: Stable; Whole exome sequencing in Suppl. Table S6 | Huntsman Cancer Institute |

**Supplementary Table S4:** Whole exome sequencing of OSUMMER.1-derived RMC-7977^R^ cell lines (Separate Excel File)

**Supplementary Table S5:** Whole exome sequencing of OSUMMER.10-derived RMC-7977^R^ cell lines (Separate Excel File)

**Supplementary Table S6:** Whole exome sequencing of PDX-4872-derived RMC-7977^R^ cell lines (Separate Excel File)

**Supplementary Table S7:** Summary of most prominent changes at the DNA and protein level in RMC-7977 resistant OSUMMER cell lines detected by Sanger sequencing or whole exome sequencing (WES) compared to parental OSUMMER.1 or OSUMMER.10 respectively.

| Cell line | Gene | DNA change | Amino Acid change |
| --- | --- | --- | --- |
| OSUMMER.1 RM1 R10nM | *Ppia* | T407G | M136R |
| OSUMMER.1 RM5 R50nM | *Map2k1* | T170G | K57T |
| OSUMMER.1 RM6 R10nM | *Map2k1* | C171G | K57N |
| OSUMMER.10 RM2 R1uM | *Ppia*<br>*Ppia* | C186G<br>C376-377AT | C62W<br>H126HX |
| OSUMMER.10 RM3 R1uM | *Map2k1* | T167G | Q56P |
| OSUMMER.10 RM4 R1uM | *Map2k1* | C362G | C121S |
| OSUMMER.1 RM6 R1uM | *Ppia* | c.100+2T>G,<br>or IVS2+2T>G<br>(568+2T>G),<br>or g.2691T>G | Splicing variant (ex2) |
| OSUMMER.10 RM1 R1uM | *Ppia* | c.3G>A (G471A) | M1I |

**Supplementary Table S8:** Composition of Mel-2 medium.

| Mel-2 medium | Final concentration |
| --- | --- |
| MCDB153 | 76% |
| L15 | 20% |
| FBS | 2% |
| Pen/Strep | 1% |
| Insulin (100x) | 1% |
| CaCl<sub>2</sub> (1.5M) | 1.68mM |
| EGF (Sigma E-4127), stock: in 0.1%BSA/HBSS | 5ng/ml |
| Bovine Pituitary Extract (BPE) | 15ng/ml |

**Supplementary Table S9:**
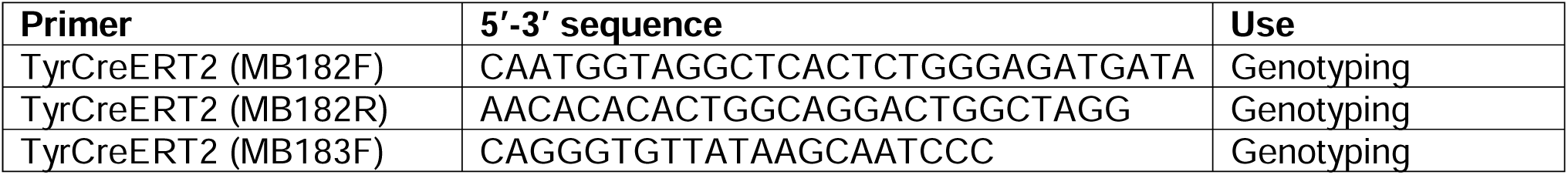
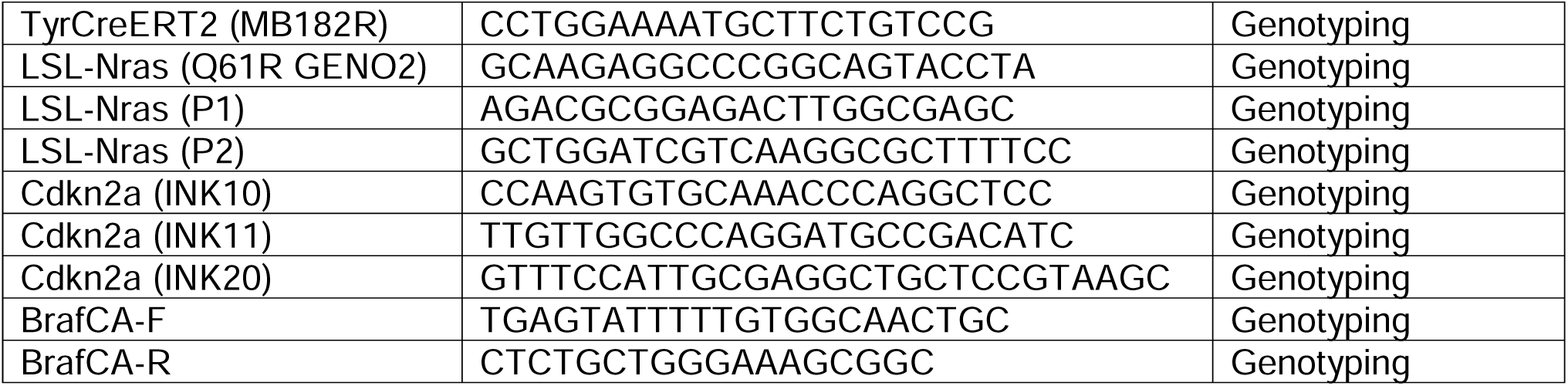
Genotyping primers.

**Supplementary Table S10.**
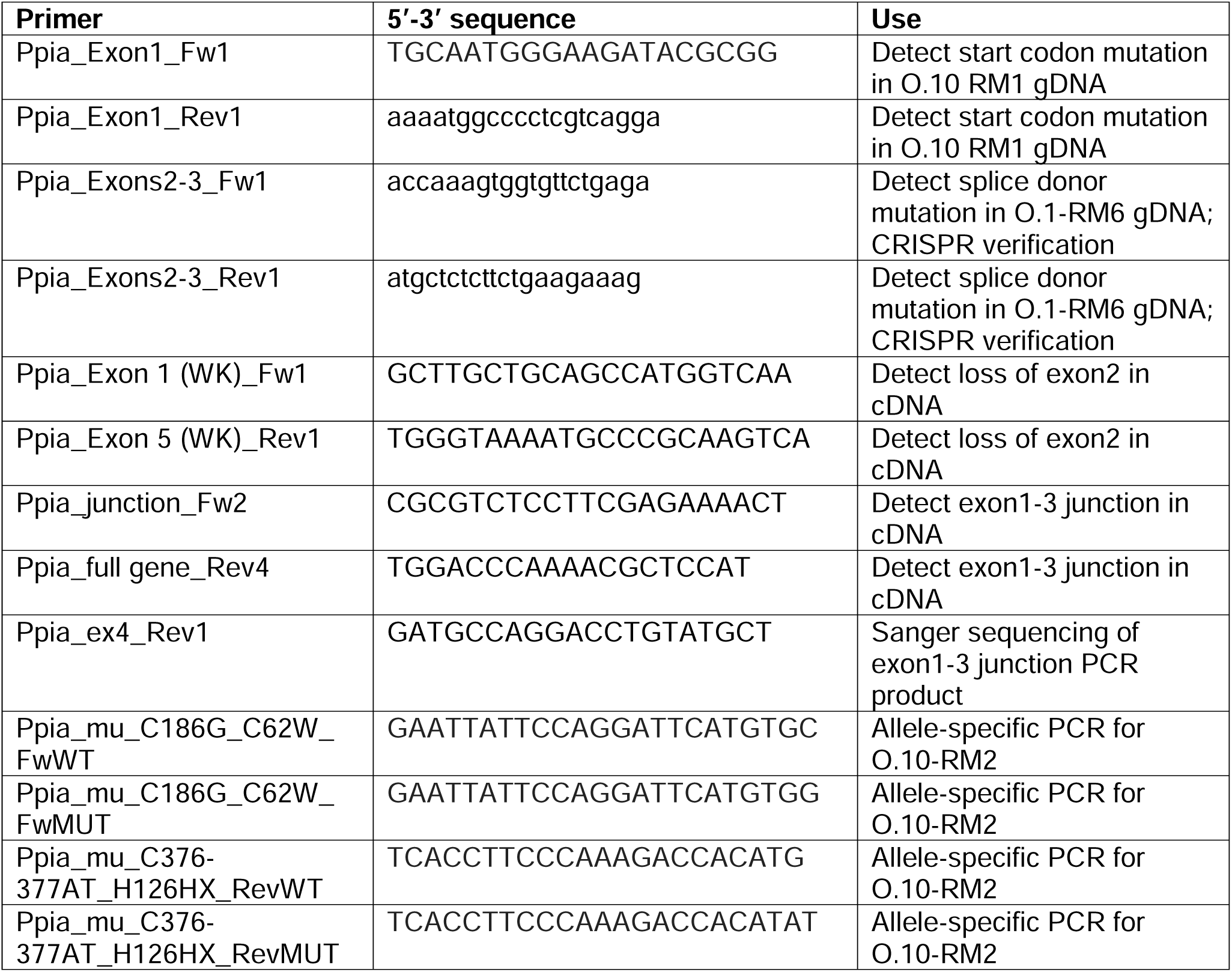
DNA Primers used for PCR or Sanger Sequencing.

**Supplementary Table S11:**
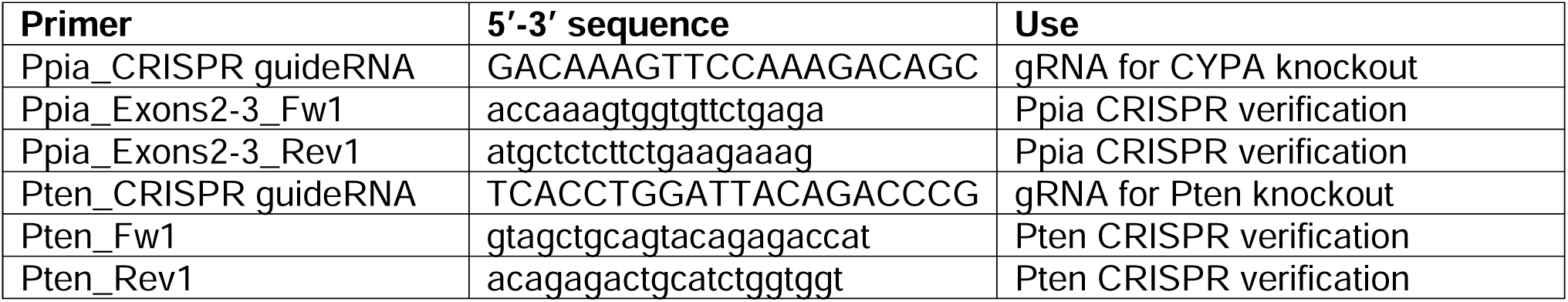
CRISPR guideRNA and associated primers.

**Supplementary Table S12:**
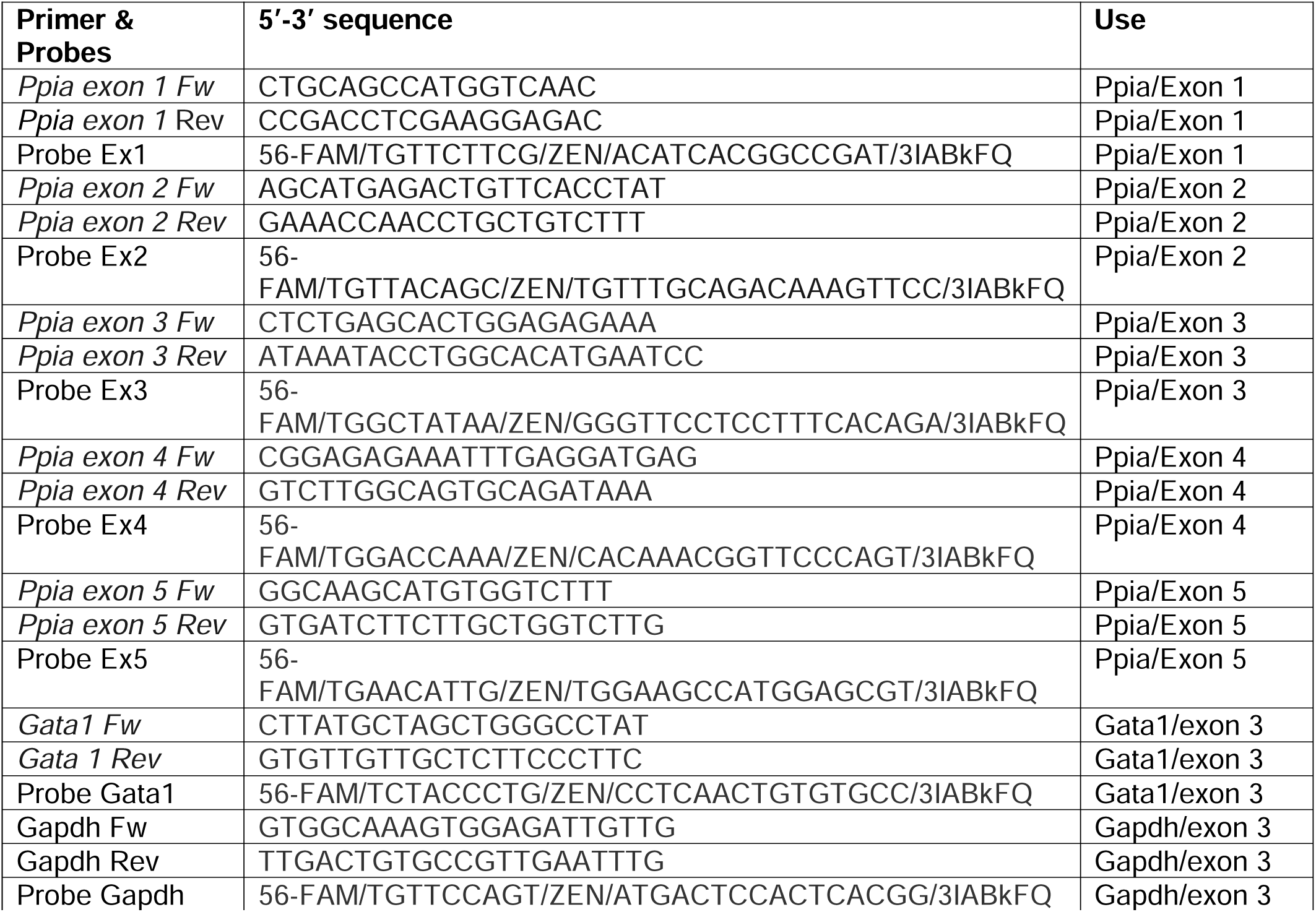
Quantitative PCR (qPCR) primers used on gDNA.

**Supplementary Table S13:**
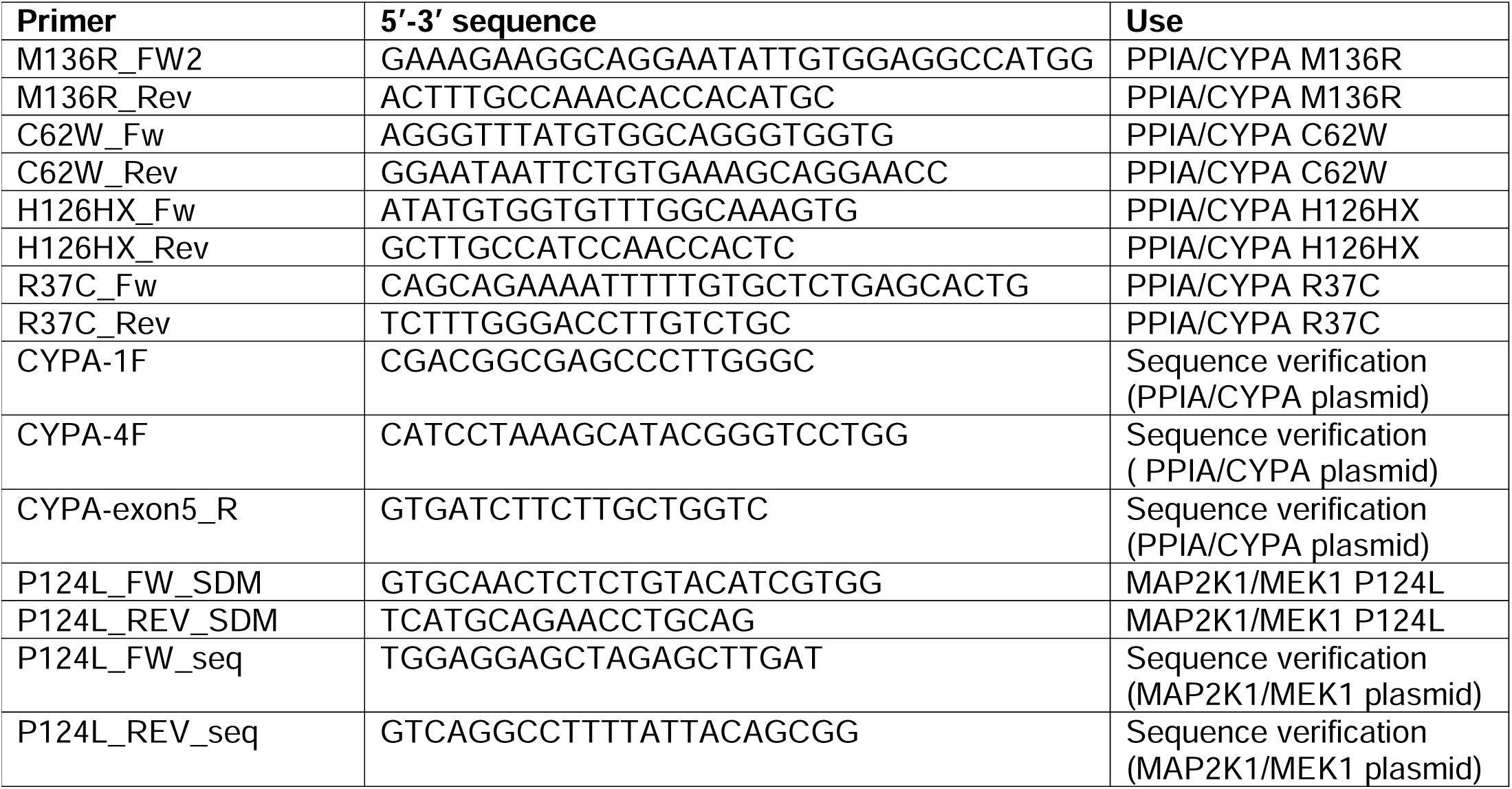
Site-directed mutagenesis primers.

